# Conjugated linolenic fatty acids trigger ferroptosis in triple-negative breast cancer

**DOI:** 10.1101/556084

**Authors:** Alexander Beatty, Tanu Singh, Yulia Y. Tyurina, Emmanuelle Nicolas, Kristen Maslar, Yan Zhou, Kathy Q. Cai, Yinfei Tan, Sebastian Doll, Marcus Conrad, Hülya Bayır, Valerian E. Kagan, Ulrike Rennefahrt, Jeffrey R. Peterson

## Abstract

Ferroptosis is a non-apoptotic form of cell death linked to the accumulation of reactive hydroperoxides generated by oxidation of polyunsaturated fatty acids (PUFAs) in membrane phospholipids. The therapeutic potential of promoting ferroptosis by enriching PUFAs in cancer cells is unknown. We found an association between elevated PUFA levels and vulnerability to ferroptosis in triple-negative breast cancer (TNBC) cells. A screen of PUFAs identified conjugated linolenic acids, including α-eleostearate, as ferroptosis inducers. Three conjugated double bonds were required for ferroptotic activity although their positioning and stereochemistry were less significant. Mechanistically, α-eleostearate differed from canonical ferroptosis inducers by a distinct dependence on acyl-CoA synthetase long-chain isoforms and by not altering glutathione or glutathione peroxidase 4 activity. Orally administered tung oil, naturally rich in α-eleostearate, limited tumor growth and metastasis in an aggressive TNBC xenograft model. These results expand our understanding of ferroptotic cell death and highlight the anti-cancer potential of conjugated PUFAs.

---

Ferroptosis is a recently described form of iron-dependent, regulated cell death that is associated with elevated lipid hydroperoxides, a form of reactive oxygen species (ROS) ^1–6^. Lipid hydroperoxides arise from enzymatic or spontaneous oxidation of membrane-associated polyunsaturated fatty acids (PUFAs). The central ferroptotic regulator is glutathione peroxidase 4 (GPX4), which is the only known peroxidase capable of efficiently reducing esterified, oxidized fatty acids into unreactive alcohols ^7^. Small molecules that trigger ferroptosis identified thus far include agents that inhibit GPX4 directly, molecules that deplete the GPX4 cofactor glutathione, and compounds that oxidize iron, all of which lead to lipid hydroperoxide accumulation ^1,8^. Elevated lipid hydroperoxides lead to disruption of membrane architecture, production of reactive aldehydes, and ultimately cell death ^9^. While the general mechanism of ferroptosis has been described, important details including metabolic features that render some cells susceptible to ferroptosis are not well understood.

Recent studies have shown that cancer cells, including renal cell carcinoma, melanoma and glioblastoma are vulnerable to ferroptosis ^10–16^. Furthermore therapy-resistant mesenchymal cancer cells exhibit a greater reliance on GPX4 activity for survival ^10,17^. These findings have highlighted the therapeutic potential of pro-ferroptotic agents, but it remains unclear whether induction of ferroptosis can be used therapeutically to selectively kill cancer cells *in vivo* ^1^. A major challenge to testing this hypothesis has been the absence of selective ferroptosis-inducing agents, such as GPX4 inhibitors, suitable for use *in vivo* ^18^.

While suppressing the GPX4-mediated antioxidant response is one approach to induce ferroptosis, a complementary approach is to promote the production of lipid hydroperoxides by increasing the availability of their PUFA precursors. Hydroperoxides can spread by a free radical-mediated chain reaction leading to oxidation of adjacent PUFAs, and consequently higher PUFA levels could increase vulnerability to the propagation of lipid peroxides. Indeed, supplementation of cells with the PUFA arachidonic acid sensitizes them to ferroptosis triggered by GPX4 inhibitors ^19^. Importantly, this approach to enhancing ferroptosis would exploit the propensity of cancer cells to scavenge fatty acids from their environment ^20^. Here we report a vulnerability in triple-negative breast cancer (TNBC) to ferroptosis associated with the accumulation of PUFAs, and identified conjugated linolenic fatty acids as PUFAs that induce ferroptosis. Mechanistically, accumulation of PUFAs in TNBC triggered ferroptosis by a mode of action distinct from canonical ferroptosis inducers. We show that oral administration of tung nut oil, naturally rich in the conjugated PUFA α-eleostearate, has anti-cancer activity in a xenograft model of TNBC. α-Eleostearate metabolites could be detected in tumors of treated mice and was associated with expression of ferroptotic markers. These results introduce a distinct class of ferroptosis inducers and offer novel insights into the molecular basis of ferroptotic sensitivity. The tractability of these dietary, pro-ferroptotic fatty acids addresses the current lack of effective GPX4 inhibitors for use *in vivo* and suggests a novel opportunity to exploit a metabolic liability in an aggressive breast cancer subtype.

## RESULTS

### Glutathione depletion triggers ferroptosis in TNBC cell lines

Survival of triple-negative breast cancer cells is dependent on the glutathione biosynthetic pathway to reduce ROS levels ^21^. The source of ROS, however, has been unclear. Treatment with buthionine sulfoximine (BSO), an inhibitor of the rate-limiting enzyme of glutathione biosynthesis, triggers cell death in BT-549 cells within days of treatment (**Fig. 1a**). Glutathione depletion in some contexts induces cell death via ferroptosis ^22^. Ferroptotic cell death is associated with increased lipid peroxidation and can be suppressed by lipophilic antioxidants like ferrostatin-1 (fer-1) and iron chelators ^1^. Consistent with ferroptosis, cytotoxicity in BSO-treated BT-549 cells was blocked by fer-1 but not by an inhibitor of apoptotic cell death, the pan-caspase inhibitor Z-VAD-FMK (**Figs. 1a, b**). Similarly, iron chelators deferoxamine (DFO) and deferiprone suppressed BSO-mediated cell death (**Fig. 1c**, Supplementary Figure S1a). Furthermore, BSO treatment resulted in the accumulation of lipid peroxidation products and this was suppressed by fer-1 (**Figs. 1d, e**). Taken together, this demonstrates that glutathione depletion triggers ferroptosis in BT-549 cells and identifies lipid hydroperoxides as the lethal source of ROS underlying glutathione addiction in TNBC cells.

**Figure 1.**
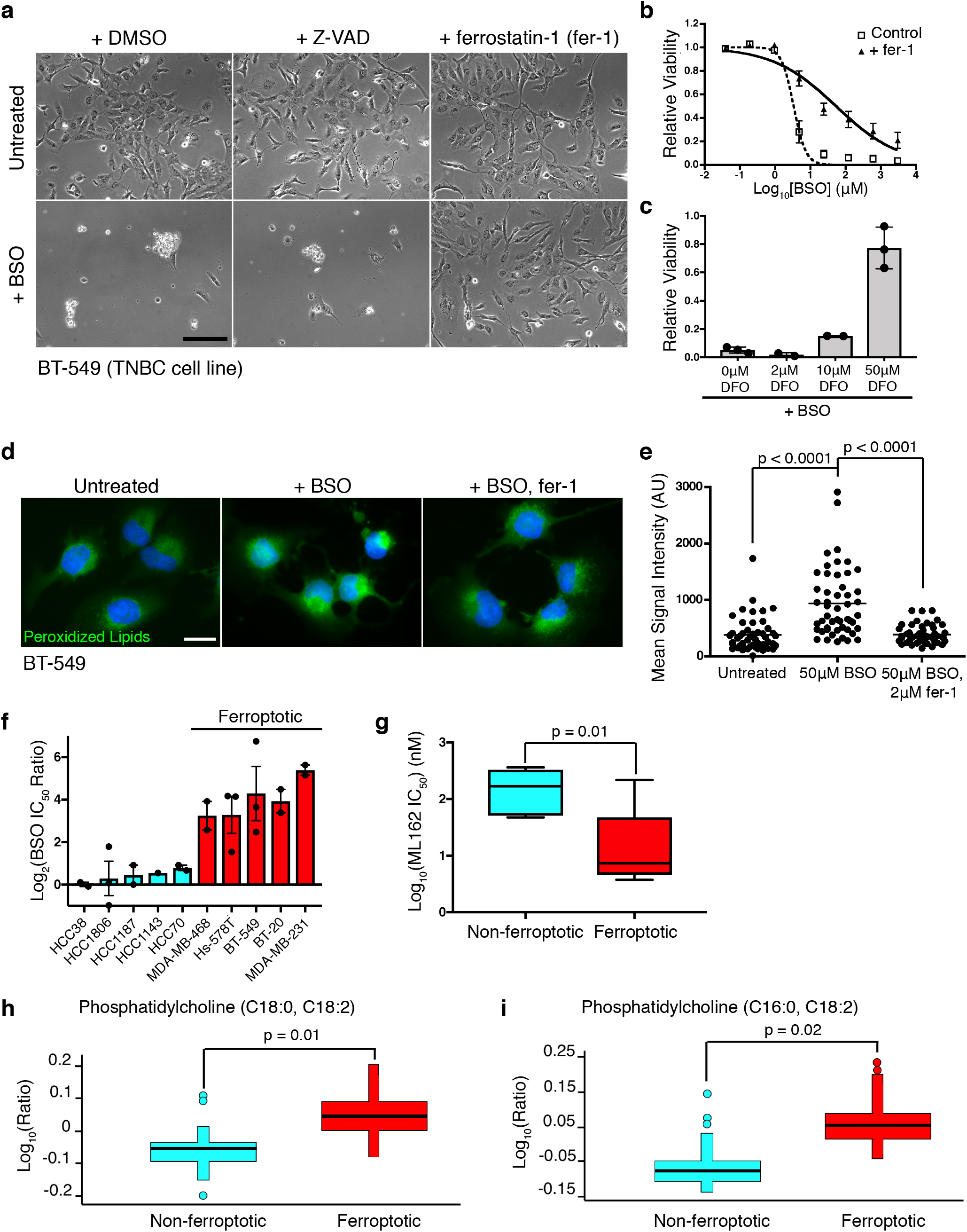
Glutathione depletion triggers ferroptosis in a subset of TNBC cell lines and elevated polyunsaturated fatty acid levels are associated with vulnerability to ferroptosis. **a**, Light micrographs of BT549 cells treated with 10 μM BSO for 72 hours in the presence or absence of 20 μM Z-VAD or fer-1. Scale bar represents 200 µm. **b**, BT-549 cell viability dose-response curves for BSO with or without 20 μM fer-1. Error bars, here and below, denote standard error of the mean. **c**, Relative viability of BT-549 cells treated with 10 μM BSO and the indicated concentration of deferoxamine (DFO) for 72 hours. Values were normalized to account for loss of viability associated with DFO. **d**, Representative fluorescent micrographs of BT-549 cells with the indicated treatment. Green corresponds to cellular macromolecules modified with breakdown products from peroxidized lipids visualized using the Click-iT lipid peroxidation detection kit (ThermoFisher Scientific). DNA is stained blue. Scale bar represents 20 µm. **e**, Quantitation of lipid peroxidation products from individual cells (n = 50 per condition) in **d**. Lines represent the mean.**f**, The log_2_-transformed ratio of the IC_50_ of BSO in the presence or absence of 2 μM fer-1 for each cell line. Cell lines in which fer-1 reduced cell death from BSO >8-fold are designated “ferroptotic”. **g**, Box and whiskers plot of the log_2_-transformed IC_50_ values for ML162 in non-ferroptotic (turquoise) and ferroptotic (red) TNBC cell lines (as defined in **f**). The line represents the median value and the whiskers show minimum and maximum values. **h**, Box plots showing the log_10_-transformed, median-normalized relative levels of phosphatidylcholine (C18:0, C18:2) and **i**, phosphatidylcholine (C16:0, C18:2) in non-ferroptotic and ferroptotic TNBC cell lines. Each cell line is represented by at least 5 replicates. The line shows the median value, the whiskers represent the upper and lower adjacent values, and outliers are shown as dots.

### Sensitivity to ferroptosis is associated with the accumulation of PUFAs

Fer-1 suppressed BSO toxicity in half of the TNBC cell lines tested (**Fig. 1f**). These “ferroptotic” cell lines (red bars) were also generally more sensitive to ML162, a small molecule inhibitor of GPX4 ^23^ (Welch’s t-test, p=0.01, **Fig. 1g**, Supplementary Figure S1b). Thus, ferroptosis is a common response to glutathione depletion in a subset of TNBC cells and is associated with a greater dependence on GPX4 activity. Notably, GPX4 transcripts were not differentially expressed between these two sets of cell lines (Supplementary Figure S1c), suggesting that ferroptotic vulnerability was independent of the level GPX4 expression. We used ANOVA of our prior metabolomic data for these same cell lines to compare levels of individual metabolites ^21^ between TNBC cell lines susceptible to BSO-mediated ferroptosis and those that are not (as defined in **Fig. 1f**), and found that two of the three most significantly differentially expressed metabolites were linoleate (18:2)-substituted phosphatidylcholines, differing only in the identity of their other acyl chain (**Fig. 1h, i**). These membrane phospholipids were more enriched in ferroptotic compared to non-ferroptotic TNBC cell lines. Because PUFAs can be directly oxidized into lipid hydroperoxides, this finding suggested the possibility that elevated PUFA-containing phospholipids may increase vulnerability to ferroptosis and underlie these cells’ greater dependence on GPX4 for survival (**Fig. 1g**). Consistent with this, recent studies implicate membrane phospholipids containing residues of arachidonic acid, and adrenic acid as promoters of ferroptosis ^19,24^.

### The conjugated PUFA α-eleostearic acid promotes cancer-selective ferroptosis

We postulated that if PUFA levels are limiting for ferroptosis, elevating levels of specific PUFAs might further sensitize TNBC cells to ferroptosis. We tested a variety of PUFA species and identified the conjugated linolenic acid isomer α-eleostearic acid [(9Z,11E,13E)-octadeca-9,11,13-trienoic acid; αESA] (**Fig. 2a**) as a fatty acid that enhanced cell death in combination with BSO (**Fig. 2b**). Unexpectedly, αESA also triggered cell death as a single agent and this death was suppressed by fer-1 (**Fig. 2b**), the iron chelator deferoxamine (**Fig. 2c)** and by the lipophilic antioxidant vitamin E (Supplementary Figure S2a) and was associated with an increase in lipid peroxidation products that could be suppressed by fer-1 (**Fig. 2d)**. Neither the pan-caspase inhibitor Z-VAD-FMK nor the necroptosis inhibitor nec-1s blocked αESA-induced cell death (Supplementary Figure S2b, c). Together, these findings demonstrate that αESA can induce ferroptosis as a single agent.

**Figure 2.**
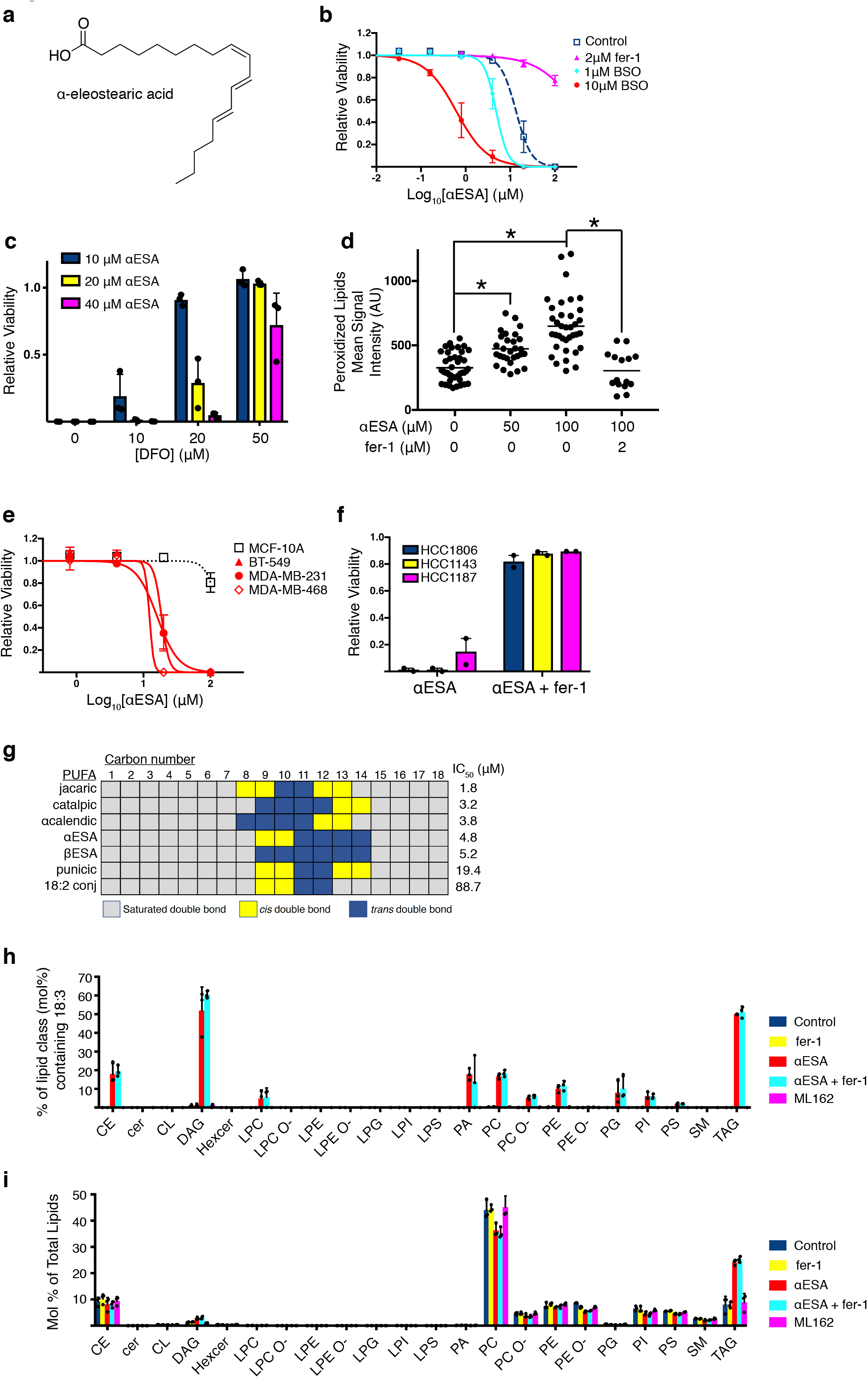
Conjugated linolenic fatty acids are incorporated into cellular lipids and induce ferroptosis in TNBC cells. **a**, Structure of α-eleostearic acid (αESA). **b**, Cell viability dose-response curves for αESA and the specified additional compound in MDA-MB-231 cells. Cells were treated for 72 hours. Error bars in this and subsequent panels represent standard error of the mean. **c**, Relative viability of MDA-MB-231 cells incubated with the indicated doses of αESA and DFO for 72 hours. Values were normalized to account for loss of viability associated with DFO. **d**, Quantitation of lipid peroxidation products in individual cells with the specified treatment. The line indicates the mean. From left to right, n = 42, 29, 36, and 14. * signifies p < 0.0001. **e**, Cell viability dose-response curves for αESA for three TNBC cell lines (red lines) and non-cancerous MCF-10A controls (black dashed line). **f**, Relative cell viability for three TNBC cell lines 72 hours after treatment with αESA or the combination of αESA and fer-1. **g**, Schematic representing the location and double bond geometry of 18-carbon conjugated polyunsaturated fatty acids. Cell viability IC_50_ values are shown for each conjugated PUFA from at least two biological replicate measurements in MDA-MB-231 cells after 72 hours treatment. **h**, Percent of the mole fraction for each of the indicated classes of lipid that contain 18:3 (18 carbon, 3 double bonds) fatty acids consistent with αESA. **i**, Mole percent of each lipid class as a fraction of total lipids in MDA-MB-231 cells incubated with vehicle, 2 μM fer-1, 50 μM αESA, 50 μM αESA and 2 μM fer-1, or 250 nM ML162 for 3 hours. Error bars in **h** and **i** show standard deviation. CE = cholesterol esters, Cer = ceramide, DAG = diacylglycerol, Hexcer = hexosylceramide, LPC = lysophosphatidylcholine, LPC O-= ether-linked lysophosphatidylcholine, LPE = lysophosphatidylethanolamine, LPE O-= ether-linked lysophosphatidylethanolamine, LPG = lysophosphatidylglycerol, LPI = lysophosphatidylinositol, LPS = lysophosphatidylserine, PA = phosphatidate, PC = phosphatidylcholine, PC O-= ether-linked phosphatidylcholine, PE = phosphatidylethanolamine, PE O-= ether-linked phosphatidylethanolamine, PG = phosphatidylglycerol, PI = phosphatidylinositol, PS = phosphatidylserine, SM = sphingomyelin, TAG = triacylglycerol

Interestingly, αESA is abundant in certain plants (e.g. bitter melon and the tung tree) ^25^ and was previously shown to suppress the growth of breast cancer cells in a manner reversible by antioxidants, though the exact mechanism was unclear ^26^. The non-transformed MCF-10A breast epithelial cell line was resistant to death by αESA (**Fig. 2e**). By contrast, all eight tested TNBC cell lines were susceptible to cell death by αESA, including cell lines that did not undergo ferroptosis in response to glutathione depletion (**Fig. 1f**), and in all cases death could be prevented by fer-1 (**Figs. 2b, f** and not shown). Time lapse phase contrast microscopy of αESA-treated BT549 cells (Supplementary Video 1) showed that during the latter part of 24 hours of treatment, otherwise healthy appearing cells undergo a sudden death, morphologically similar to cell death by the canonical ferroptosis inducer ML162 (Supplementary Video 2) and distinct from apoptotic death induced by staurosporine (Supplementary Video 3). Cell death by αESA was associated with a large membrane protrusion that ultimately ruptured, accompanied by a cessation of cytoplasmic motility.

### Structure-activity analysis of conjugated PUFAs

αESA is a naturally occurring fatty acid with no structural similarity to known ferroptosis inducers. To characterize structural features of αESA required for ferroptosis, we screened structurally related conjugated and unconjugated PUFAs for death induction in BT549 cells. αESA possesses conjugated double bonds at carbons 9, 11, and 13 with cis, trans, trans stereochemistry, respectively (**Fig. 2g**). We examined cis-trans stereoisomers of αESA and found that isomerization at position 9 (β–eleostearic acid) or at both 9 and 13 positions (catalpic acid) retained cell death activity while isomerization at position 13 alone (punicic acid) led to an ~4-fold reduction in potency. Jacaric acid (conjugated 18:3, cis-8, trans-10, cis-12) showed the most potent ability to kill cells (1.8 µM IC_50_), demonstrating that shifting the positioning of the double bonds while maintaining the sequential cis, trans, cis stereochemistry of punicic acid, did not disrupt activity. α–Calendic acid (conjugated 18:3, trans-8, trans-10, cis-12), a stereoisomer of jacaric acid, was similarly potent. Thus, cell killing is not strictly dependent on the precise positioning or stereochemistry of double bonds within the fatty acid. A conjugated linoleic acid (18:2), which shares similar position and stereochemistry of its two double bonds with αESA, was a poor inducer of cell death, however, suggesting that third conjugated double bond is required for its activity. Furthermore, all tested non-conjugated PUFAs (e.g. docosahexaenoic, linoleic, and linolenic acids) were less potent single-agent inducers of cell death (Supplementary Figure S2d) including arachidonic acid (20:4), which was of interest because it enhances ferroptosis in some contexts ^19,27^. Cell killing by arachidonic acid could not be rescued by fer-1 (Supplementary Figure S2e). By contrast, jacaric- and catalpic acid-induced death, like that caused by αESA, was suppressed by fer-1 (Supplementary Figure S2f). Similar results were found in BT-549 cells (Supplementary Figure S2g), demonstrating the conserved ability of diverse conjugated 18:3 fatty acids to trigger ferroptosis.

### αESA is incorporated into diverse cellular lipids

αESA, unlike canonical ferroptotic agents, is a fatty acid that could potentially be incorporated as an acyl chain in cellular lipids. To examine how αESA is metabolized, we conducted a mass spectrometry-based lipidomic analysis of MDA-MB-231 cells that were untreated, αESA-treated, fer-1-treated, or treated with both αESA and fer-1 for three hours. While 18:3-containing lipids were extremely rare in untreated cells, αESA-treatment was associated with incorporation of 18:3 acyl chains, into a diverse array of lipid species including phospholipids, cholesterol esters and storage lipids and fer-1 did not dramatically alter the spectrum of lipid classes that incorporated 18:3 acyl chains (**Fig. 2h**; detailed results are presented in Supplementary Table 1). The abundance of di- and triacylglycerol lipids increased ~2-fold in αESA- and αESA + fer-1-treated cells compared to untreated controls, suggesting that a portion of αESA is incorporated into storage lipids (**Fig. 2i**). All lipid classes significantly altered by αESA treatment were similarly altered by αESA in the presence of fer-1 (Supplementary Table 1). αESA-induced lipidomic changes were not simply downstream consequences of ferroptosis since the effects were not replicated by ML162 treatment (**Fig. 2h, i**). Together these results demonstrate that αESA is incorporated into cellular fatty acid pools in a largely fer-1-insensitive manner.

### Conjugated PUFAs exhibit anti-cancer activity and promote expression of ferroptotic markers *in vivo*

We used tung oil as an inexpensive and rich source of αESA (80% of all fatty acids in tung oil are αESA ^28^) to assess whether its anti-TNBC activity could be recapitulated in the aggressive MDA-MB-231 orthotopic xenograft model. Once tumors reached ~ 75 mm^3^, mice were treated by oral gavage five times per week with 100 µl tung oil. Control mice received high-oleic (18:1) safflower oil. Body weights of tung oil-treated mice did not differ significantly from controls over the treatment period (not shown). However, tung oil significantly suppressed tumor growth (**Fig. 3a**) and endpoint tumor weight (**Fig. 3b**) and also substantially reduced lung metastatic invasion compared to controls (**Figs. 3c, d**). Immunohistochemistry revealed a 10% decrease in Ki67-positive cells in tumors from tung-treated mice compared to controls and a modest, though statistically significant increase in tumor cleaved caspase 3-positive cells, a marker of apoptosis (from 2% to 5%) (Supplementary Figure S3a, b).

**Figure 3.**
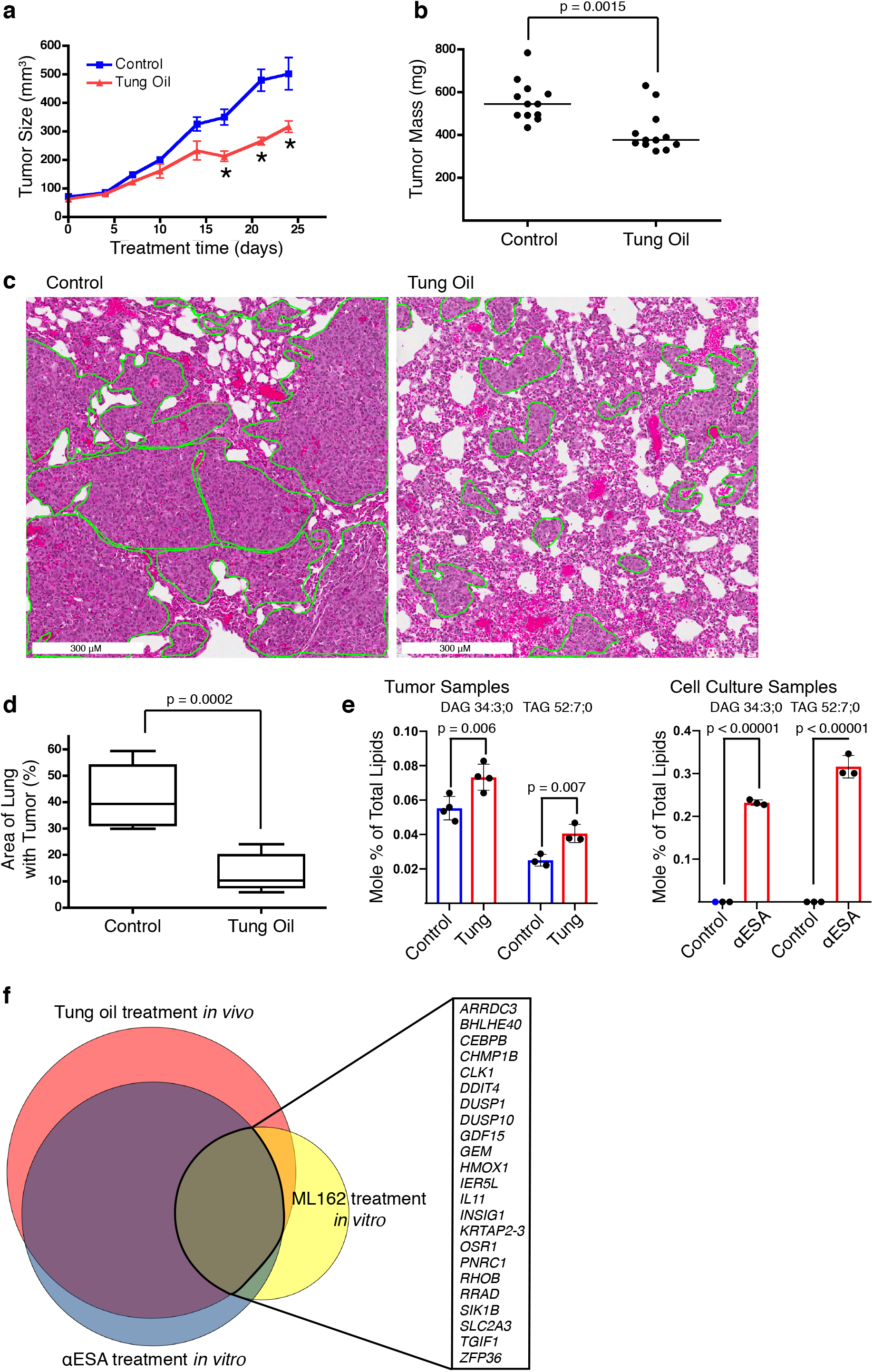
Tung oil suppresses TNBC xenograft growth and metastasis with markers of ferroptosis. **a**, Tumor volumes over time for orthotopic MDA-MB-231 xenografts in NSG mice treated orally with either safflower oil (control) or tung oil (n = 6 mice per group with bilateral tumors). * denote p values, which are 0.003, 0.00001, and 0.0003 from left to right. Error bars here and below show standard error of the mean. **b**, Final tumor masses of individual tumors following 24 days of treatment. Lines represent median values. **c**, Representative hematoxylin and eosin-stained lung sections. Regions containing metastatic TNBC cells are outlined. **d**, Quantification of the percentage of lung area infiltrated by metastatic cells. The line represents the median value and the whiskers show minimum and maximum values. **e**, Mole percent of total lipids for DAG 34:3 and TAG 52:7 in tumors treated with tung oil (left) or in MDA-MB-231 cells grown in culture and incubated with 50 μM αESA for 3 hours (right). The p values were calculated using a one-sided Student’s t-test, not adjusted for multiple hypothesis testing. **f**, Proportional Venn diagram showing the overlap in genes with transcript levels that were significantly altered by > 2-fold (0.05 false discovery rate) after a 5-hour incubation with 100 μM αESA or 125 nM ML-162 or in orthotopic MDA-MB-231 xenograft tumors treated as described above for **a**. Genes commonly altered in all three data sets are listed.

To provide evidence that the reduction in tumor growth is associated with tumor exposure to αESA, we conducted a lipidomic analysis of αESA-treated and control tumors. The two most differentially expressed lipids between the tumor groups were diacylglycerol (34:3) and triacylglycerol (52:7), which were increased in tung oil-treated tumors by 30% and 60%, respectively (**Fig. 3e**). The masses of these lipids are consistent with at least one linolenate moiety, although their precise acyl chain composition was not determined. Notably, the expression of these two lipids was also increased in cells treated with αESA *in vitro*, compared to untreated cells (**Fig. 3e**). Our findings identify the neutral lipids diacylglycerol (34:3) and triacylglycerol (52:7) as potential biomarkers of tumor exposure to αESA. Further, they suggest that circulating αESA, released from tung oil, is taken up by TNBC tumor cells and incorporated into lipids.

Next, we assessed whether tumors in αESA-treated mice expressed RNA markers of ferroptosis. We conducted RNA sequencing of three biological replicates each of tumors from treated and control mice and compared the differentially expressed genes with a separate *in vitro* experiment using cultured MDA-MB-231 cells treated with either αESA, the GPX4 inhibitor ML162, or corresponding vehicle controls. Using a false discovery rate of 5% and a filter of >2-fold change in RNA expression level, 124 genes were altered in tung oil-treated tumors compared to safflower oil-treated controls (**Fig. 3f**). Significantly, 89 genes (89/122, 73%) altered in tung oil-treated tumors were also altered in αESA-treated cells in culture. Fisher’s exact test showed extremely significant overlap between these gene sets (p=3.2 × 10^−246^) and all 89 overlapping genes were altered in the same direction (Supplementary Table 2). This is consistent with our hypothesis that the inhibition of tumor growth *in vivo* is related to the cell death caused by αESA *in vitro*. Importantly, there was also a highly significant overlap in genes altered by tung oil *in vivo*, αESA *in vitro*, and by *in vitro* treatment with ML162 (**Fig. 3f** and Supplementary Table 2). Fisher’s exact test between any two gene sets demonstrated p-values of < 7.2 × 10^−61^. Furthermore, all of the shared genes were altered in the same direction. These results demonstrate highly concordant gene signatures between tung oil treatment and αESA treatment, consistent with the hypothesis that tumors in tung oil-treated mice are exposed to αESA. Furthermore, the highly significant overlap in the signatures of αESA- and ML162-treated cells provide strong evidence that both treatments trigger a similar response.

All three treatments altered a common set of 23 genes (**Fig. 3f** and Supplementary Table 2) including genes previously identified as increased during ferroptosis including RHOB, SLC2A3, DDIT4 and HMOX1 ^15,29^. In addition, the previously reported ferroptosis marker CHAC1 ^29^ was commonly upregulated in both tung-oil treated tumors and αESA-treated cells. 66 genes were altered by both αESA and tung oil but not ML162 (Supplementary Table 2), suggesting a potential source of biomarkers for αESA-induced ferroptosis.

### Ferroptosis induction by αESA is not mediated by GPX4 inhibition

Next, we sought mechanistic insights into ferroptosis induction by αESA. One possibility is that αESA could promote the formation of lipid hydroperoxides by inhibiting GPX4 either directly or indirectly. For example, ML162 and the structurally distinct GPX4 inhibitor RSL3 directly bind and inactivate GPX4 while BSO and erastin indirectly inhibit GPX4 by starving GPX4 of its essential cofactor glutathione ^1^. While BSO treatment suppressed glutathione levels, treatment of cells with αESA for 24 hours did not significantly affect total cellular glutathione levels despite decreasing cell viability (**Fig. 4a**), suggesting that αESA does not trigger ferroptosis by depleting glutathione.

**Figure 4.**
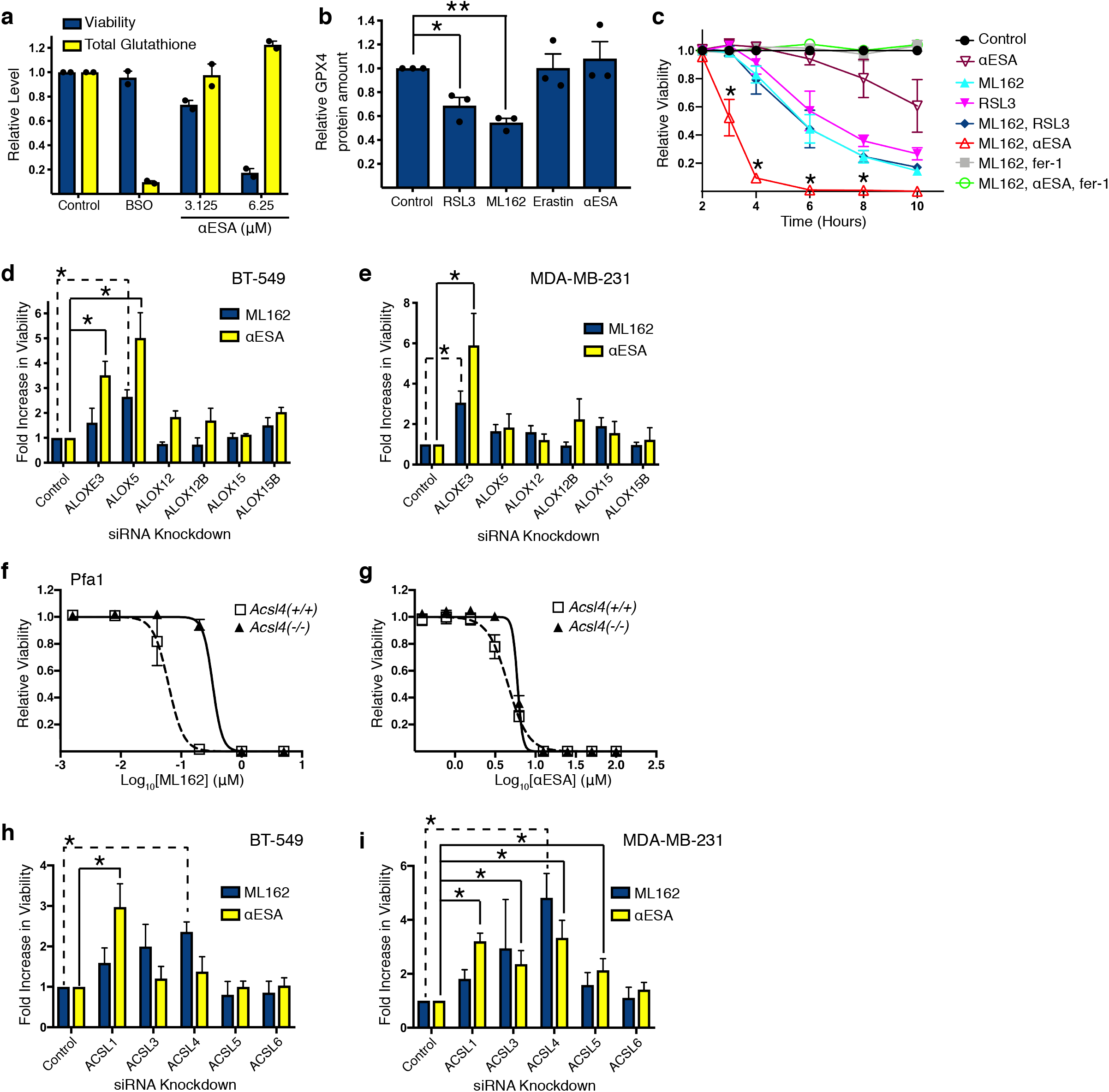
Mechanistic analysis of αESA-induced ferroptosis. **a**, Relative viability (blue) and levels of total glutathione (yellow) in MDA-MB-231 cells after 24 hours of treatment with 20 μM BSO, or the specified dose of αESA. Error bars in this and subsequent panels represent standard error of the mean. **b**, Densitometric quantification of GPX4 protein levels from western blots of lysates from MDA-MB-231 cells that were treated for 4 hours with the indicated agent. * denotes p = 0.01, and ** indicates p = 0.0002, n = 3 independent experiments. **c**, Cell viability over time for MDA-MB-231 cells treated as specified. * denote p values from n ≥ 3 independent experiments. p values from left to right are 0.003, 0.0001, 0.02, and 0.008. Bar chart showing fold-change in relative cell viability after 72-hour siRNA knockdown of the specified lipoxygenase followed by 24-hour treatment with either ML-162 (blue) or αESA (yellow) in **d**, BT-549 (n = 3) or **e**, MDA-MB-231 cells (n ≥ 4). * denote p values below the significance threshold and greater than 2-fold suppression of cell toxicity. The p values for BT-549 from left to right are 0.004, 0.01, and 0.02. The p values for MDA-MB-231 are 0.01 and 0.01. Cell viability dose-response curves for control and Acsl4-deficient Pfa1 cells treated with **f**, ML-162 or **g**, αESA. **h**, Bar charts showing fold change in the fraction of viable BT-549 (n ≥ 3) or **i**, MDA-MB-231 cells (n ≥ 3), relative to cells transfected with a non-targeting siRNA, 72 hours after transfection with the designated ACSL siRNA followed by 24-hour treatment with either ML-162 (blue) or αESA (yellow). * signify p values below the significance threshold and greater than 2-fold suppression of cell toxicity. The p values from left to right for BT-549 are 0.005 and 0.01 and for MDA-MB-231 are 0.005, 0.0003, 0.03, 0.03, and 0.04. Complete data for the bar charts in **d**, **e**, **h**, and **i** can be found in Supplementary Table S3.

Direct GPX4 inhibitors have been shown to reduce GPX4 protein levels ^19^. We confirmed this for ML162 and RSL3 but found that αESA or the negative control erastin had no effect on GPX4 levels (**Fig. 4b**), suggesting that αESA does not inhibit GPX4 activity in a similar way. As an alternative approach to determine whether αESA inhibits GPX4 either directly or indirectly, we compared the kinetics of ferroptosis induction in TNBC cells treated with αESA, ML162 and RSL3 alone and in combination. The combination of both GPX4 inhibitors at 500 nM triggered fer-1-suppressible cell death with similar kinetics to either GPX4 inhibitor alone, indicating this dose of inhibitor was sufficient to achieve near maximal GPX4 inhibition (**Fig 4c**). In contrast, cell viability was lost significantly more rapidly when cells were co-treated with 500 nM ML162 and 50 µM αESA, a dose of αESA that took longer to induce cell death than either of the GPX4 inhibitors alone. Consistent with ferroptosis, cell death triggered by the combination of ML162 and αESA could be fully rescued by fer-1 (**Fig 4c**). The ability of αESA to further enhance the kinetics of cell death under conditions of maximal GPX4 inhibition suggests that αESA and GPX4 inhibitors trigger ferroptosis by distinct mechanisms.

### αESA-induced ferroptosis is lipoxygenase and ACSL1-dependent

Ferroptosis is intimately associated with PUFA peroxidation ^30,31^. Lipoxygenases have been reported to mediate peroxidation in ferroptosis induced by GPX4 inhibition ^24,27^ but non-enzymatic lipid oxidation by Fenton chemistry may also contribute to this process ^18,32^. To assess lipoxygenase contributions to αESA-induced ferroptosis we systematically knocked down lipoxygenase isoforms by RNA interference in two TNBC cell lines. ALOXE3 knockdown most significantly suppressed αESA toxicity in both cell lines (**Fig. 4d, e**), demonstrating a role for this lipoxygenase isoform in αESA-induced death. This finding does not, however, rule out a potential role for non-enzymatic oxidation, since αESA undergoes spontaneous oxidation more rapidly than other unconjugated and conjugated linolenic acids ^33,34^.

The first step in esterification of fatty acids into cellular lipids is their conjugation to CoA, catalyzed by enzymes of the acyl-CoA synthetase family. There are five long chain-specific isoforms (ACSLs) and loss of ACSL4 significantly protects cells from ferroptosis induced by direct or indirect inhibition of GPX4 (**Fig. 4f**; ^16,35,36^). αESA, however, was similarly toxic to ACSL4-deficient Pfa1 mouse embryonic fibroblast cells and control parental cells (**Fig. 4g**), suggesting a difference in the mechanism of action of αESA and canonical ferroptosis inducers. αESA-induced cell death was rescued by fer-1 treatment in both cell lines, consistent with ferroptosis (Supplementary Figure S4a). To determine if any ACSL isoforms contribute to ferroptosis induced by αESA, we individually knocked down expression of each of the five ACSL isoforms in BT549 and MDA-MB-231 cells by RNAi. Knockdown of individual ACSL isoforms expressed in each cell line was confirmed by quantitative PCR (Supplementary Figures S4b, c). As expected, ML162 toxicity was most effectively suppressed by ACSL4 depletion in both cell lines. αESA toxicity, however, was most significantly suppressed by knockdown of ACSL1 (**Figs. 4h, i**). Thus, αESA differs in its ACSL isoform dependence compared to a conventional ferroptosis inducer, further supporting a mechanism distinct from canonical ferroptosis inducers.

### αESA triggers ACSL1-dependent increases in lipid hydroperoxides

As ferroptosis is associated with an accumulation of oxidized phospholipids, we next used liquid chromatography–mass spectrometry (LC–MS) to ask if αESA treatment impacts phospholipid oxidation. We quantified 141 mono-, di-, and tri-oxygenated lipids across seven lipid classes in BT-549 cells and found the abundance of oxidized lipids was significantly increased in cells treated with αESA (p = 0.02) (**Fig. 5a**). Overall, the amounts of 51 (36%) lipid species were significantly changed and a majority of those were increased (41/51, 80%). The classes with both the highest number and frequency of increased oxidized species were cardiolipins (17/36 47%) and phosphatidylethanolamines (18/40 45%) and included two di-oxygenated phosphatidylethanolamine species containing arachidonate residues (**Fig. 5b**) that have been previously identified as ferroptotic signals in cells treated with the GPX4 inhibitor RSL3 ^19^. These results are consistent with our observation of increased lipid peroxidation products in αESA-treated cells (**Fig. 2d**) and suggest that oxidized PE and CL phospholipids may contribute to ferroptosis triggered by αESA.

**Figure 5.**
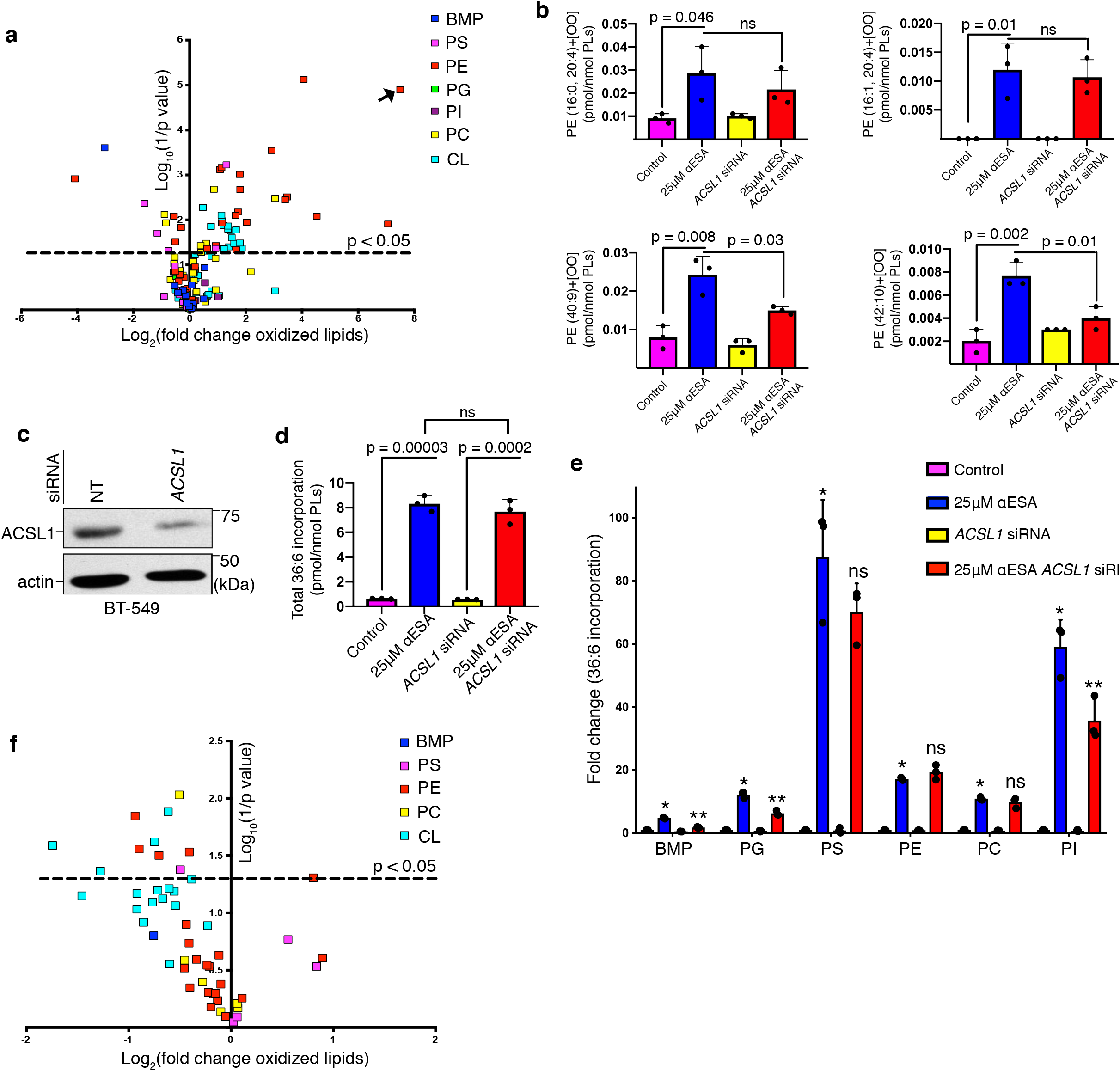
αESA treatment increases cellular oxidized phospholipids and a subset of them are decreased by ACSL1 depletion. **a**, Volcano plot showing log_2_(fold change) and significance (log_10_(1/p)) for oxidized phospholipids in BT-549 cells incubated for 8 hours with 25 μM αESA compared to cell incubated with vehicle (methanol). Each lipid species is colored according to phospholipid class. Abbreviations are as follows: BMP = bis(monoacylglycero)phosphate, PS = phosphatidylserine, PE = phosphatidylethanolamine, PG = phosphatidylglycerol, PI = phosphatidylinositol, PC = phosphatidylcholine, CL = cardiolipin. The arrow indicates a lipid species for which a fold change could not be computed because it was only detected after αESA treatment. **b**, Bar charts showing the amounts of di-oxygenated PE species that were significantly increased by αESA treatment compared to vehicle-treated controls. **c**, Western blot showing ASCL1 protein in BT-549 cells 72 hours after transfection with either non-targeting siRNA or *ACSL1* siRNA. β-actin is the loading control. Relative amount of all phospholipids quantified that contain 2 acyl chains with 36 carbons and 6 double bonds (36:6, consistent with αESA incorporation) in **d**, all phospholipids quantified or **e**, individual phospholipid classes in BT-549 cells 72 hours after transfection with either non-targeting siRNA or *ACSL1* siRNA and then treated with either vehicle or 25 μM αESA for 8 hours. In **e**, * indicate significant changes compared to control (p ≤ 0.001) and ** denote significant changes compared to αESA-treated cells transfected with non-targeting siRNA (significant p values from left to right are 0.00004, 0.001, 0.02). **f**, Volcano plot showing log_2_(fold change) versus log_10_(1/p-value) for oxidized phospholipids in BT-549 cells transfected with *ACSL1* siRNA and incubated with 25 μM αESA for 8 hours compared to αESA-treated cells transfected with non-targeting siRNA. The set is limited to the 51 oxidized phospholipid species that were significantly changed αESA treatment as shown in **a**.

One possible explanation for the ACSL1-dependence of αESA-induced ferroptosis is that ACSL1 mediates the incorporation of αESA into cellular lipids. To test this, we sought to quantify the presence of 18:3 fatty acids in diverse lipid species in αESA- and control-treated cells in which ACSL1 expression was silenced (**Fig. 5c**). Total acyl chain length and double bond count were characterized for each lipid species containing two acyl chains (excludes cardolipins) and we focused on 36:6 lipids because this composition is consistent with two αESA acyl chains. Validating this approach, αESA treatment led to a 13-fold increase in total 36:6 lipids (**Fig. 5d**), which were distributed across all examined lipid classes (**Fig. 5e**). Silencing ACSL1 did not affect overall 36:6 lipid levels in αESA-treated cells, demonstrating that ACSL1 is not essential for αESA esterification, although some individual classes exhibited significantly decreased 36:6 lipids. An alternative hypothesis is that ACSL1 depletion suppresses αESA-induced ferroptosis by lessening the accumulation of oxidized lipid species. We quantified oxygenated lipids in αESA-treated cells in which ACSL1 was knocked down (**Fig. 5f**). Of the oxygenated lipids that were significantly increased following incubation with αESA, 10 (10/41, 24%) were responsive to ACSL1 depletion and all were decreased (detailed in Supplementary Figure S5), consistent with the idea that decreasing ACSL1 levels may reduce the accrual of specific peroxidized phospholipids that induce ferroptosis in αESA-treated cells. Indeed, we found that some, but not all, αESA-induced, di-oxygenated PE lipids were decreased on ACSL1-silencing (**Fig. 5b**).

### Small-molecule enhancers of αESA-induced ferroptosis implicate phospholipase A2 as a suppressor of ferroptosis

Since αESA causes an accumulation of lipid hydroperoxides (**Fig. 5c**) without decreasing GPX4 (**Fig. 4b**), its toxicity might be enhanced in combination with GPX4 inhibitors. Consistent with this, low doses of BSO, which on their own do not affect cellular viability, enhanced αESA toxicity *in vitro* (**Fig. 2b**). Similarly, treatment of mice with BSO at a dose that does not affect tumor growth on its own ^21^, enhanced the anti-tumor activity of tung oil in MDA-MB-231 orthotopic xenografts (**Fig. 6a**). We likewise found that subtoxic doses of αESA enhanced cell killing by the GPX4 inhibitors ML162 and RSL3 in multiple cell lines (**Fig. 6b** and Supplementary Figure S6). Taken together, our observations are consistent with αESA promoting an increase in lipid hydroperoxides that is opposed by, but ultimately overwhelms GPX4 activity. In addition, they suggest that combining GPX4 inhibitors, should these become clinically viable, with αESA may enhance the anti-tumor activity of αESA.

**Figure 6.**
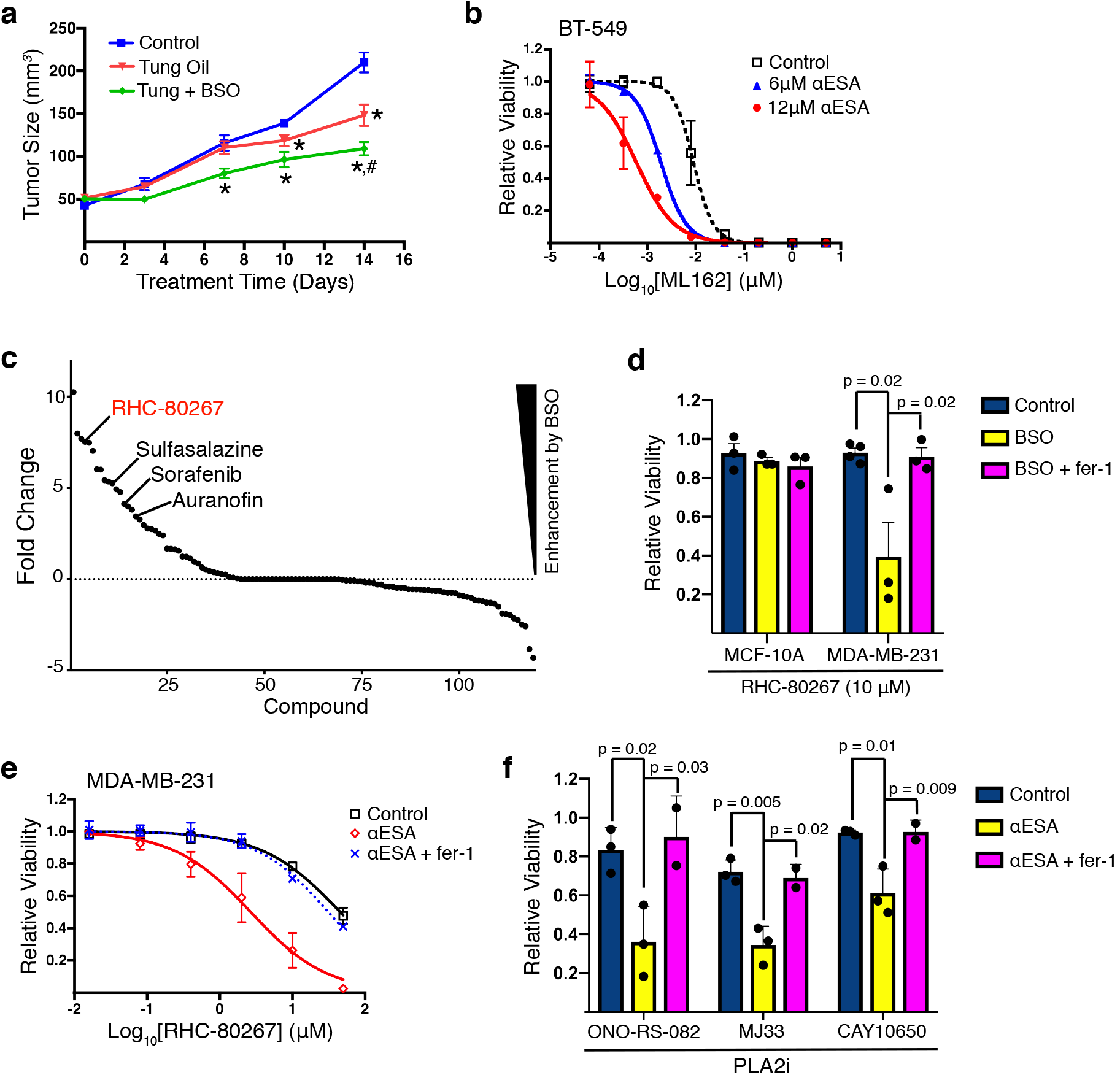
Identification of compounds that enhance ferroptosis by αESA. **a**, Tumor volumes for orthotopic MDA-MB-231 xenograft tumors in NSG mice treated for 14 days by oral gavage with either 100 µl safflower oil (control, n = 6 mice, bilateral tumors), tung oil (n =4), or tung oil and BSO given at 20 mM in drinking water (n = 4). * denote p < 0.05 compared to the control. The p values from left to right are 0.02 and 0.006 for tung oil-treated mice, and 0.009, 0.0001, and 0.000003 for tung oil and BSO-treated mice. # indicates p = 0.004 compared to tung oil-treated mice. Error bars in this and subsequent panels show standard error of the mean. **b**, Cell viability dose-response curves for ML162 and the indicated dose of αESA in BT-549 cells after 72 hours of treatment. **c**, The log_2_-transformed ratio of the IC_50_ of compounds included in a collection of small molecule modulators of cellular metabolism (Focus Biomolecules) in the presence or absence of 10 μM BSO in MDA-MB-231 cells treated for 72 hours. **d**, Relative cell viability in MDA-MB-231 or MCF-10A cells incubated with RHC-80267 alone, with a sensitizing dose of BSO, or with BSO and fer-1. **e**, Cell viability dose-response curves for RHC-80267 alone, with a minimally lethal, sensitizing dose of αESA (3-6 µM), or with both αESA and fer-1 in MDA-MB-231 cells. p values generated to assess rescue of cell viability by fer-1 were calculated using a one-sided Student’s t-test. **f**, Relative cell viability in MDA-MB-231 and the specified PLA2 inhibitors (ONO-RS-082 40 μM, MJ33 5 μM, CAY10650 20 μM) with vehicle, a sensitizing dose of αESA or with αESA and fer-1.

Finally, we sought to identify other metabolic pathways that might modulate ferroptosis in TNBC cells. We conducted a multi-dose screen of 126 known small molecule inhibitors of diverse metabolic pathways for their ability to modify the toxicity of glutathione depletion in MDA-MB-231 cells. Both enhancers and suppressors of BSO toxicity were identified, including previously reported enhancers of ferroptosis, auranofin ^37, 22^, and sorafenib ^29^ (**Fig. 6c**). Among the most potent enhancers was the multi-targeting compound RHC-80267. As a single agent, RHC-80267 only weakly induced cell death (**Fig. 6d, e**). In combination with subtoxic BSO, however, RHC-80267 induced fer-1 suppressible cell death, demonstrating a synthetic lethal interaction between these two compounds (**Fig. 6e**). In untransformed MCF-10A cells BSO did not enhance RHC-80267 toxicity (**Fig. 6d**). RHC-80267-induced cell death was also enhanced by αESA in multiple cell types and could be fully rescued by fer-1 (**Fig. 6e;** other cell lines not shown). Thus, RHC-80267 is a selective enhancer of ferroptosis triggered by either αESA or canonical ferroptosis inducers.

One target of RHC-80267 is diacylglycerol lipase ^22^ but knock down of DAGL1 and DAGL2 isoforms individually and together did not affect αESA potency (not shown). RHC-80267 also exhibits inhibitory activity against phospholipase A2 (PLA2) ^38^. To probe a potential role for PLA2 isoforms we used three additional, structurally unrelated PLA2 inhibitors, each of which enhanced ferroptotic death by αESA (**Figs. 6f**). In every case, cell death was suppressed by fer-1. These results point to a key role for PLA2 in protecting against both BSO- and αESA-induced ferroptosis. PLA2 isoforms play a role in phospholipid repair by liberating oxidatively damaged fatty acids from membrane phospholipids ^39^. Thus, inhibition of PLA2 may promote αESA-induced ferroptosis by allowing the accumulation of oxidized fatty acids in membrane phospholipids.

## DISCUSSION

Cancer cells, particularly drug-resistant and metastatic cells are vulnerable to oxidative stress and depend on antioxidant pathways for their survival ^10,40-45^. This metabolic liability could potentially be exploited by two complementary therapeutic approaches, inhibiting anti-oxidant survival pathways or promoting oxidative stress. Consistent with the first strategy, we and others have demonstrated a dependence on glutathione-based anti-oxidant pathways in breast cancer ^21,42,46,47^. Here, we demonstrate the feasibility of the second strategy, enhancing oxidative stress using conjugated linolenic acids to trigger the production of reactive hydroperoxides, a key mediator of ferroptosis.

GPX4 acts as a central regulator of ferroptosis by preventing the accumulation of phospholipid hydroperoxides. Existing small-molecule inducers of ferroptosis directly or indirectly inhibit GPX4 activity, leading to the accumulation of lipid peroxides produced endogenously. We show here that conjugated linolenic fatty acids provide a distinct mechanism to initiate ferroptosis in which these oxidizable fatty acids increase the production of lipid peroxides that overwhelm GPX4 activity. Furthermore, the combination of these two approaches robustly enhances cell death both *in vitro* and *in vivo* (**Fig. 4c, Figs. 6a, b**). Supporting a distinct mechanism of action, the pro-ferroptotic activity of αESA relies on a subset of ACSL and lipoxygenase isoforms that is at least partially distinct from those exploited by canonical ferroptosis inducers. Despite these mechanistic differences, αESA and conventional ferroptosis inducers both lead ultimately to an increase in lipid peroxides (**Figs. 1e, 2d**) and induce a similar transcriptional response (**Fig. 3f**), suggesting a convergent mechanism of lethality. While GPX4 inhibition allows the accumulation of endogenous lipid hydroperoxides, αESA is incorporated into membrane lipids, increasing levels of oxidizable PUFAs, and enhancing overall lipid oxygenation. GPX4 deletion in adult mice leads to acute renal failure and death ^48^, suggesting that the therapeutic window for clinical GPX4 inhibitors, should they become available, may be narrow. By contrast, tung oil and αESA (unpublished data) appear to be well tolerated in mice. Thus, these findings expand our understanding of the pathways that regulate ferroptosis and provide a new approach for exploiting ferroptosis therapeutically.

Our metabolic inhibitor screen uncovered a potential role for PLA2 in preventing ferroptosis (**Fig. 6**), consistent with a recent study that identified secreted PLA2 as a suppressor of lipotoxic stress in breast cancer cells ^49^. Inhibitors of PLA2 potentiate ferroptosis by both αESA (**Fig. 6f**) and canonical ferroptosis inducers (**Fig. 6d**), possibly by preventing oxidatively damaged fatty acyl chains from being liberated from membrane lipids ^39^. This finding suggests that PLA2 inhibitors might enhance the therapeutic efficacy of conjugated linolenic acids or canonical ferroptosis inducers in cancer. PLA2 isoforms constitute a large family of enzymes with potential functional redundancy, making it challenging to identify whether specific isoforms protect against ferroptosis. Nevertheless, this finding supports an emerging model in which dynamic remodeling of membrane fatty acids can modulate cellular vulnerability to PUFA oxidation ^50^.

There are two major limitations to translating ferroptosis into a therapeutic modality, the lack of orally available ferroptosis inducers and the absence of robust biomarkers of ferroptosis for use *in vivo*. We discovered that tung oil, a naturally rich source of αESA, suppresses xenograft tumor growth and metastasis and provide evidence that this activity is mediated by ferroptosis. These results identify tung oil as an inexpensive and bioavailable agent potentially suitable for clinical ferroptosis induction. Moreover, its lack of toxicity and positive interactions with other pro-ferroptotic agents, reported here, suggest opportunities for combination therapy. Finally, we identified a set of genes whose expression is altered in cells undergoing αESA- and tung oil-induced ferroptosis as potential biomarkers of ferroptosis and a source of candidate genes that may contribute to this cell death mechanism.

## ACKNOWLEDGEMENTS

This work was supported in part by NIH GM083025, U19AI068021, HL114453-06, CA165065-06, NS076511, the PA Breast Cancer Coalition, the Fifth District AHEPA Cancer Research Foundation, the Spurlino Family Foundation, the Rita Hollman Foundation, the Eileen Stein Jacoby Fund, and Translation Research and In Vino Vita awards from Fox Chase Cancer Center. This work was supported by NIH CORE Grant P30 CA006927. We thank Drs. S. Balachandran and J. Karanicolas for comments on the manuscript.

## METHODS

### Cell lines

MDA-MB-468 cells were obtained by the Cell Culture Facility at Fox Chase Cancer Center from ATCC as part of the NCI-60 panel. All other TNBC cell lines and the MCF10A cell line were obtained directly from the American Type Culture Collection (ATCC, Manassas, VA 20110, USA). The suppliers routinely authenticate the cell lines by short tandem repeat profiling though only MDA-MB-231 cell lines were authenticated by our laboratory (April 2018). All cell lines were amplified and frozen within 2 months of receipt. TNBC cell lines were cultured in RPMI-1640, 10% heat inactivated fetal bovine serum, 2 mM supplemental glutamine, and 100 μg/mL penicillin/streptomycin with the exception of BT-20, which was cultured in MEM with the same supplements. MCF-10A cells were cultured in high calcium medium with 5% horse serum as described ^51^. Mouse embryonic fibroblast (Pfa1) *Acsl4*-deficient cells were described previously ^16^. Cells were cultured in a humidified incubator at 37 °C with 5% CO_2_. Cell lines are periodically tested for *Mycoplasma* contamination using DAPI (4′,6-diamidino-2-phenylindole) to stain DNA.

### Statistical Analysis

Metabolite profiling data shown in Figure 1h and i were log_10_-transformed before further analysis to achieve an approximate normal distribution. Missing values were not imputed for univariate analysis. Mixed model analysis of variance (ANOVA) using R with package nlme was applied to identify differentially expressed intracellular metabolites in non-ferroptotic (HCC38, HCC1806, HCC1143, HCC70) compared to ferroptotic (MDA-MB-468, MDA-MB-231, BT-549, BT-20, Hs-578T) TNBC cell lines. Our previously reported data set of 155 metabolites including 5-6 replicates per cell line ^21^ was taken considering “cell line” as random factor and “ferroptosis response” as categoric factor. ANOVA models was read out concerning t-statistics results comprising estimates, t-values, and p-values. Significance level was set to an α-error of 5%. The multiple test problem was addressed by calculating the false discovery rate (FDR) using the Benjamini & Hochberg method. In general, other statistical comparisons were performed using Student’s t-test unless noted and the threshold for significance was p < 0.05. Data is reported as mean and standard error of the mean of at least 2-3 independent experiments unless otherwise stated. All data points reflect measurements of distinct samples.

### Lipidomic analysis

For Figures 2h and i, triplicate samples of MDA-MB-231 cells at ~ 60% confluence were incubated with vehicle (methanol), 2 μM fer-1, 50 μM αESA, 50 μM αESA and 2 μM fer-1, or 250 nM ML162 for 3 hours and then were typsinized, washed twice with phosphate buffered saline (PBS) without magnesium or calcium, and resuspended at a concentration of 1500 cells/μL, and frozen in liquid nitrogen. For tumor samples (Figure 3e), four biological replicate tumor samples per condition were homogenized in PBS without magnesium or calcium using a dounce homogenizer to yield a sample containing 5 mg of tumor (wet weight) / mL and were then frozen in liquid nitrogen.

Mass spectrometry (MS)-based lipid analysis in Figures 2 and 3e were performed by Lipotype GmbH (Dresden, Germany) as described ^52^. Lipids were extracted using a two-step chloroform/methanol procedure ^53^. Samples were spiked with internal lipid standard mixture containing: cardiolipin 16:1/15:0/15:0/15:0 (CL), ceramide 18:1;2/17:0 (Cer), diacylglycerol 17:0/17:0 (DAG), hexosylceramide 18:1;2/12:0 (HexCer), lyso-phosphatidate 17:0 (LPA), lyso-phosphatidylcholine 12:0 (LPC), lyso-phosphatidylethanolamine 17:1 (LPE), lyso-phosphatidylglycerol 17:1 (LPG), lyso-phosphatidylinositol 17:1 (LPI), lyso-phosphatidylserine 17:1 (LPS), phosphatidate 17:0/17:0 (PA), phosphatidylcholine 17:0/17:0 (PC), phosphatidylethanolamine 17:0/17:0 (PE), phosphatidylglycerol 17:0/17:0 (PG), phosphatidylinositol 16:0/16:0 (PI), phosphatidylserine 17:0/17:0 (PS), cholesterol ester 20:0 (CE), sphingomyelin 18:1;2/12:0;0 (SM), triacylglycerol 17:0/17:0/17:0 (TAG). After extraction, the organic phase was transferred to an infusion plate and dried in a speed vacuum concentrator. The dried extract was re-suspended in 7.5 mM ammonium acetate in chloroform/methanol/propanol (1:2:4, V:V:V) and the second step dry extract was re-suspended in a 33% ethanol solution of methylamine in chloroform/methanol (0.003:5:1; V:V:V). All liquid handling steps were performed using the Hamilton Robotics STARlet robotic platform. Samples were analyzed by direct infusion on a QExactive mass spectrometer (Thermo Scientific) equipped with a TriVersa NanoMate ion source (Advion Biosciences). Samples were analyzed in both positive and negative ion modes with a resolution of Rm/z=200=280000 for MS and Rm/z=200=17500 for MS-MS experiments, in a single acquisition. MS-MS was triggered by an inclusion list encompassing corresponding MS mass ranges scanned in 1 Da increments (Surma et al. 2015). Both MS and MS-MS data were combined to monitor CE, DAG and TAG ions as ammonium adducts; PC, PC O-, as acetate adducts; and CL, PA, PE, PE O-, PG, PI and PS as deprotonated anions. MS only was used to monitor LPA, LPE, LPE O-, LPI and LPS as deprotonated anions; Cer, HexCer, SM, LPC and LPC O-as acetate adducts. Data were analyzed using lipid identification software based on LipidXplorer ^54,55^. Data post-processing and normalization were performed using an in-house developed data management system (Lipotype). Only lipid identifications with a signal-to-noise ratio >5, and a signal intensity 5-fold higher than in corresponding blank samples were considered for further data analysis.

### Live Cell Imaging

For the time lapse imaging, BT-549 cells were seeded in a 6-well plate at 125,000 cells/well. After 24 hours, stauroporine (500 nM), αESA (25 μM), or ML162 (500 nM) were added along with 2 mM HEPES (pH 7.4) and then each well was overlaid with mineral oil and placed in a temperature-controlled imaging chamber (37 °C). Three fields in each well were imaged every 10 minutes for a 24-hour period using a Nikon Eclipse TE300 inverted microscope equipped with a Nikon Plan Fluor 10x NA 0.30 objective and a QImaging Retiga EXi camera and using MetaVue software. Movies were assembled using the Fiji implementation of ImageJ ^56^. For photomicrographs, cell images were taken using a Nikon Eclipse TE2000 inverted microscope equipped with Nikon Plan Fluor 10x NA 0.30 objective using Metavue (MetaMorph) or Ocular (QImaging) software. The micrographs shown in Figure S2b were captured using the EVOS XL Core Imaging System (ThermoFisher Scientific) with the 10x objective.

### Detection of lipid peroxidation products

The Click-iT Lipid Peroxidation Imaging kit (ThermoFisher Scientific) with Alexa Fluor 488 azide was used to detect macromolecules modified with breakdown products from peroxidized lipids according to the recommendations of the manufacturer. Cells were incubated with linoleamide alkyne (LAA) reagent for 16 hours in experiments involving BSO and 4 hours for experiments involving αESA before being fixed with fixed with 4% formaldehyde. DAPI was used to visualize DNA. Fluorescent micrographs were captured with a Nikon Eclipse TE2000 inverted microscope with a Nikon Plan Apo 60x A/1.40 Oil using Metavue (MetaMorph) or Ocular (QImaging) software. The fluorescence signal intensity of individual cells was quantified using ImageJ (NIH).

### Small interfering RNA

For siRNA knockdown experiments, cells were transfected (DharmaFECT 1, ThermoFisher Scientific) with ON_TARGETplus SMART pools (ThermoFisher Scientific) targeting individual *ACSL* and *ALOX* family members, *DAGL1*, *DAGL2*, or a non-targeting siRNA pool (Pool #1, D-001206-13, ThermoFisher Scientific). Efficiency of mRNA depletion was assessed 72 hours post-transfection using qPCR or western blot in the case of *ACSL1* (Proteintech, 13989-1-AP). In experiments to determine suppression of cell toxicity resulting from αESA or ML162 by silencing expression of individual *ACSL* or *ALOX* genes, doses of αESA or ML162 were selected such that there was 5-15% remaining viability in cells transfected with non-targeting siRNA following 24 hours of treatment.

### Quantification of cellular phospholipids

BT-549 cells were transfected with *ACSL1* or non-targeting siRNA 72 hours prior to incubation with 25 μM αESA or vehicle (methanol) for 8 hours. After 8 hours, cells (~ 4-6 × 10^6^ per replicate) were trypsinized, collected, washed once with phosphate-buffered saline, and frozen in liquid nitrogen. Triplicate samples were collected for each condition. MS analysis of PLs was performed on an OrbitrapTM FusionTM LumosTM mass spectrometer (ThermoFisher Scientific). Phospholipids were separated on a normal phase column (Luna 3 µm Silica (2) 100Å, 150 × 2.0 mm, (Phenomenex)) at a flow rate of 0.2 mL/min on a Dionex Ultimate 3000 HPLC system. The column was maintained at 35 °C. The analysis was performed using gradient solvents (A and B) containing 10 mM ammonium formate. Solvent A contained propanol:hexane:water (285:215:5, v/v/v) and solvent B contained propanol:hexane:water (285:215:40, v/v/v). All solvents were LC/MS grade. The column was eluted for 0-23 min with a linear gradient from 10 % to 32 % B; 23-32 min using a linear gradient of 32-65% B; 32-35 min with a linear gradient of 65-100 % B; 35-62 min held at 100% B; 62-64 min with a linear gradient from 100 % to 10 % B followed by and equilibration from 64 to 80 min at 10 % B. The instrument was operated with ESI probe in negative polarity mode. Analysis of LC/MS data was performed using software package Compound Discoverer (ThermoFisher Scientific) with an in-house generated analysis workflow and oxidized phospholipid database.

### Quantitative PCR

Total RNA was isolated using an RNeasy kit (Qiagen) and tested for quality on a Bioanalyzer (Agilent Technologies). RNA concentrations were determined with a NanoDrop spectrophotometer (ThermoFisher Scientific). RNA was reverse transcribed using Moloney murine leukemia virus reverse transcriptase (Ambion-ThermoFisher Scientific) and a mixture of anchored oligo-dT and random decamers (Integrated DNA Technologies). Two reverse-transcription reactions were performed for each sample using either 100 or 25 ng of input RNA. Aliquots of the cDNA were used to measure the expression levels of the genes with the primers, and Power SYBR Green master mix (Applied Biosystems, ThermoFisher Scientific) on a 7900 HT sequence detection system (Applied Biosystems, ThermoFisher Scientific). Cycling conditions were 95°C, 15 min, followed by 40 (two-step) cycles (95°C, 15 s; 60°C, 60 s). Ct (cycle threshold) values were converted to quantities (in arbitrary units) using a standard curve (four points, four-fold dilutions) established with a calibrator sample. The primers used were as follows: ACSL1 (GACATTGGAAAATGGTTACCAAATG, GGCTCACTTCGCATGTAGATA), ACSL3 (CGAAGCTGCTATTTCAGCAAG, CTGTCACCAGACCAGTTTCA), ACSL4 (TCTTCTCCGCTTACACTCTCT, CTTATAAATTCTATCCATGATTTCCGGA), ACSL5 (GGAGAATACATTGCACCAGAGA, ACTCCTACTAAGGATGACCGT), and GPX4 (ACGTCAAATTCGATATGTTCAGC, AAGTTCCACTTGATGGCATTTC). 36B4 was used as the normalizer (CCCATTCTATCATCAACGGGTACAA, CAGCAAGTGGGAAGGTGTAATCC).

### RNAseq

Total RNA was isolated using the RNeasy kit (Qiagen) from three independent replicates of MDA-MB-231 cells incubated for 5 hours with ML162 (125 nM) and αESA (100 μM) and 2 replicates for the control incubated with the vehicle for αESA (methanol). Three biological replicate tung-oil or control MDA-MB-231 xenograft tumor samples were homogenized in ice-cold PBS using a dounce homogenizer and total RNA was isolated using the RNeasy kit (Qiagen). The stranded mRNA-seq library was generated from 1000 ng of total RNA from each sample using Truseq stranded mRNA library kit (Illumina) according to the product instructions. In short, mRNAs were enriched twice via poly-T-based RNA purification beads, and subjected to fragmentation at 94 °C for 8 minutes via the divalent cation method. The first strand cDNA was synthesized by SuperscriptII (ThermoFisher Scientific) and random primers at 42 °C for 15 mins, followed by second strand synthesis at 16 °C for 1 hour. During second strand synthesis, the dUTP was used to replace dTTP, thereby the second strand was quenched during amplification. A single ‘A’ nucleotide is added to the 3’ ends of the blunt fragments at 37 °C for 30 minutes. Adapters with illuminaP5, P7 sequences as well as indices were ligated to the cDNA fragment at 30 °C for 10 minutes. After Ampure bead (BD) purification (Beckman Coulter), a 15-cycle PCR reaction was used to enrich the fragments. PCR was set at 98 °C for 10 s, 60 °C for 30 s and extended at 72 °C for 30 s. Libraries were again purified using AmPure beads, checked for quality check with a Bioanalyzer (Agilent Technologies) and quantified with Qubit (Invitrogen). Sample libraries were subsequently pooled and loaded to the HiSeq2500 and sequenced using a Hiseq rapid SRcluster kit and HiSeq rapid SBS kit (Illumina). Single 50bp reads were generated for the bioinformatic analysis. Raw sequence reads were aligned to the human genome (hg38) using the Tophat algorithm ^57^ and Cufflinks algorithm ^58^ was implemented to assemble transcripts and estimate their abundance. Cuffdiff ^59^ was used to statistically assess expression changes in quantified genes in different conditions.

### Cell viability assays and reagents

Cells were seeded in 96-well plates (Corning 3917, 3125-6250 cells per well) and treated with compounds 24 hours after plating. Compounds were purchased from Cayman Chemical with the exception of linolenic acid, linoleic acid, docosahexaenoic acid, conjugated linoleic acid (16413), arachidonic acid, Z-VAD-FML, ONO-RS-083, and (+)-α-tocopherol (T3634), which were purchased from Sigma Aldrich. RSL3 was purchased from Selleckchem, catalpic acid, α-calendic acid, and β-ESA were obtained from Larodan Fine Chemicals, and punicic acid was purchased from Matreya LLC. Initial aliquots of ferrostatin-1, deferoxamine, and ML162 were generous gifts of the Dixon Lab (Stanford) and additional quantities were purchased from Cayman Chemical. Cell viability was measured using CellTiter-Glo Luminescent Cell Viability Assay (Promega) according to the manufacturer’s instructions. Luminescence was measured on an EnSpire Alpha (Perkin Elmer) using the integrated software package. Data were normalized to vehicle-treated or sensitizing agent-alone controls and IC_50_ curves were produced with GraphPad Prism (Version 8).

### Glutathione measurements

Cellular glutathione was quantified using the GSH/GSSG-Glo kit (Promega) according to the instructions provided by the manufacturer. Drug-treated samples were normalized to parallel cell viability measurements using the CellTiter-Glo assay (Promega).

### Quantitation of GPX4 protein levels

MDA-MB-231 were seeded at a density of 170,000 cells/well of a 6-well plate. After 24 hours, Cells were incubated for 4 hours with either vehicle (methanol or DMSO) ML162 (200 nM), RSL3 (200nM), erastin (1 μM), or αESA (50 μM). Following treatments, cells were typsinized, pelleted, lysed with RIPA buffer. The western blot was performed using a GPX4 antibody (Abcam, [EPNCIR144], ab125066) (1:1000), β-actin antibody (Abcam, ab8227) (1:5000) for loading control, and goat anti-rabbit IgG (H+L) secondary antibody, HRP (ThermoFisher Scientific, 31460) (1:3000). ImageJ (NIH) was used for the densitometric quantification.

### Screen of metabolic modulators

A cell viability screen was performed using a library of 126 small molecule modulators of metabolic pathways (Focus Biomolecules IntelliScreen Cellular Metabolism Library). MDA-MB-231 cells were plated in 384-well format at a density of 1250 cells per well in the presence or absence of 10 μM BSO. Compounds (5 doses per compound) were delivered by pin transfer 24 hours after cell plating. After 72 hours, cell viability was measured using CellTiter-Glo Luminescent Cell Viability Assay (Promega) according to the manufacturer’s instructions. Data were normalized to vehicle-treated controls and were fitted to IC_50_ curves using GraphPad Prism (Version 8).

### Mouse xenografts

Orthotopic xenografts were generated by implanting 2.5 million MDA-MB-231 cells in 100 µL phosphate-buffered saline (PBS) mixed with 100 µL growth factor-reduced Matrigel (Corning) bilaterally into the fourth inguinal fat pad of four- to six-week-old female NOD.*Cg-Prkdc^scid^Il2rg^tm1Wjl^*/SzJ (NSG) mice. After 14 days, when tumors were roughly 50 mm^3^, animals were randomized into treatment groups. Mice were treated on weekdays with either safflower oil (Whole Foods 365) and Tung oil (Sigma-Aldrich 440337) at a dose of 100 µL administered by oral gavage. L-BSO (Sigma) was administered via the drinking water (20 mM) *ad libitum* as previously reported ^21,60^. The Fox Chase Cancer Center Institutional Animal Care and Use Committee reviewed and approved all animal procedures as compliant with relevant ethical regulations.

### Tissue preparation, histology, and immunohistochemistry

Tumor sections and lung sections were fixed in 10% formalin for 24-48 hours, dehydrated and embedded in paraffin. Immunohistochemical staining was performed by standard methods. Tissue sections were incubated with primary antibodies to anti-human Ki-67 (Clone MIB-1, DAKO) and cleaved caspase-3 (Cell Signaling; #9661) at 4 °C in a humidified chamber overnight. To quantify lung metastases, lungs from three mice per treatment group were fixed, sectioned and hematoxylin and eosin stained. Ten equally sized regions (2 from each lobe) were randomly selected and scored manually for area occupied by cancerous tissue. Micrographs were captured with a Nikon DS-Fi1 camera (Melville, NY, USA) on a Nikon Eclipse 50i microscope and quantified using a ScanScope CS5 slide scanner (Aperio).

### Data Availability

All data generated or analyzed during this study are included in this published article (and its supplementary information files) or are available from the corresponding author on reasonable request.

