## Supplementary Figures for "Conjugated linolenic fatty acids trigger ferroptosis in triple-negative breast cancer"

#### SUPPLEMENTARY FIGURE LEGENDS

**Figure S1. A subset of TNBC cells undergo ferroptosis following glutathione depletion and are sensitive to GPX inhibition. a**, Relative viability of BT-549 cells incubated with 10 μM BSO and the specified dose of deferiprone for 72 hours. Values were normalized to account for loss of viability associated with deferiprone. Error bars in this and the subsequent panel show standard error of the mean. **b**, Cell viability dose response curves for ML162 in 10 TNBC cells lines and the non-transformed mammary epithelial cell line MCF-10A. **c**, Relative mRNA expression of GPX4 measured by qPCR in the indicated cell lines. In **b** and **c**, non-ferroptotic and ferroptotic TNBC cell lines (as defined in **Fig 1f)** are colored turquoise and red, respectively.

**Figure S2. Conjugated linolenic acids trigger ferroptosis in TNBC cells. a**, Relative viability dose response curves for αESA in BT-549 cells co-treated with the specified dose of vitamin E (α-tocopherol) for 48 hours. Error bars denote standard deviation. **b**, Light micrographs of BT-549 cells treated with 25 μM αESA for 48 hours in the presence or absence of 20 μM Z-VAD or 2 μM fer-1. **c**, Relative viability of BT-549 cells incubated for 72 hours with either vehicle or 20 μM αESA and either 20 μM Z-VAD, 50 μM nec-1s, or 2 μM fer-1. Errors bars show standard error of the mean. **d**, $Log_2(IC_{50})$ values for the indicated polyunsaturated fatty acid in MDA-MB-231 after 72 hours of incubation. Error bars show standard error of the mean. **e**, Cell viability dose response curves for arachidonic acid in BT-549 after 72 hours of treatment with or without 2 μM fer-1. Error bars show standard deviation. Cell viability dose response curves for the noted conjugated linolenic acid in the presence or absence of 2 μM fer-1 in **f**, MDA-MB-231 or **g**, BT-549 cells. Treatment time was 48 hours for BT-549 and 72 hours for MDA-MB-231. Error bars indicate standard error of the mean.

**Figure S3. Markers for apoptosis and cell proliferation in tung oil-treated xenograft tumors.** Bar charts showing the percent of cells in orthotopic MDA-MB-231 xenograft tumors from mice treated with safflower oil (control) or tung oil that were positive for **a**, cleaved caspase-3 or **b**, Ki67. Error bars denote standard error of the mean.

**Figure S4. Mechanism of ferroptosis by αESA. a**, Cell viability dose response curves for αESA in control and *Acsl4*-deficient mouse embryonic fibroblast (Pfa1) cells with or without 2 μM fer-1. Error bars show standard deviation. Relative mRNA expression of ACSL family members by qPCR in **b**, BT-549 and **c**, MDA-MB-231 72 hours after transfection with the indicated *ACSL* siRNA. Error bars depict standard error of the mean. BT-549 do not express *ACSL5* and neither BT-549 nor MDA-MB-231 express *ACSL6* at a level that could be quantified.

**Figure S5. Quantitative LC/MS assessment of oxidized phospholipids altered by αESA treatment in BT-549 cells.** Radar plot showing significant increases in the amount of **a**, mono- and **b**, di-oxygenated PE species after treatment with 25 μM αESA for 8 hours. Amount of phospholipid (pmol/μmol of total phospholipid) is shown in magenta for controls and blue for αESA-treated cells for all radar plots. **c**, Bar charts showing the amounts of PE species that were significantly increased by αESA treatment compared to vehicle-treated controls and decreased in αESA-treated cells in which ACSL1 had been depleted compared to αESA-treated cells. Data are based on three replicates and error bars show standard deviation. The color scheme shown in this panel applies to subsequent bar charts. **d**, Radar plot showing significant increases in oxidized CL species after treatment with αESA. **e**, Bar charts showing the amounts of oxidized CL species that were significantly increased by αESA treatment and responsive to ACSL1 knockdown. **f**, Radar plot showing significant increases in oxidized PC species after treatment with αESA. Amounts of **g**, a PC species and **h**, a PS species that were significantly increased by αESA treatment and decreased by ACSL1 depletion.

**Figure S6. Ferroptosis triggered by GPX4 inhibition is enhanced by αESA in TNBC cells.** Relative cell viability dose response curves for ML162 in MDA-MB-231 cells and vehicle, 3 μM αESA, or 2 μM fer-1. Cells were treated for 72 hours and error bars show standard error of the mean.

#### Figure S1

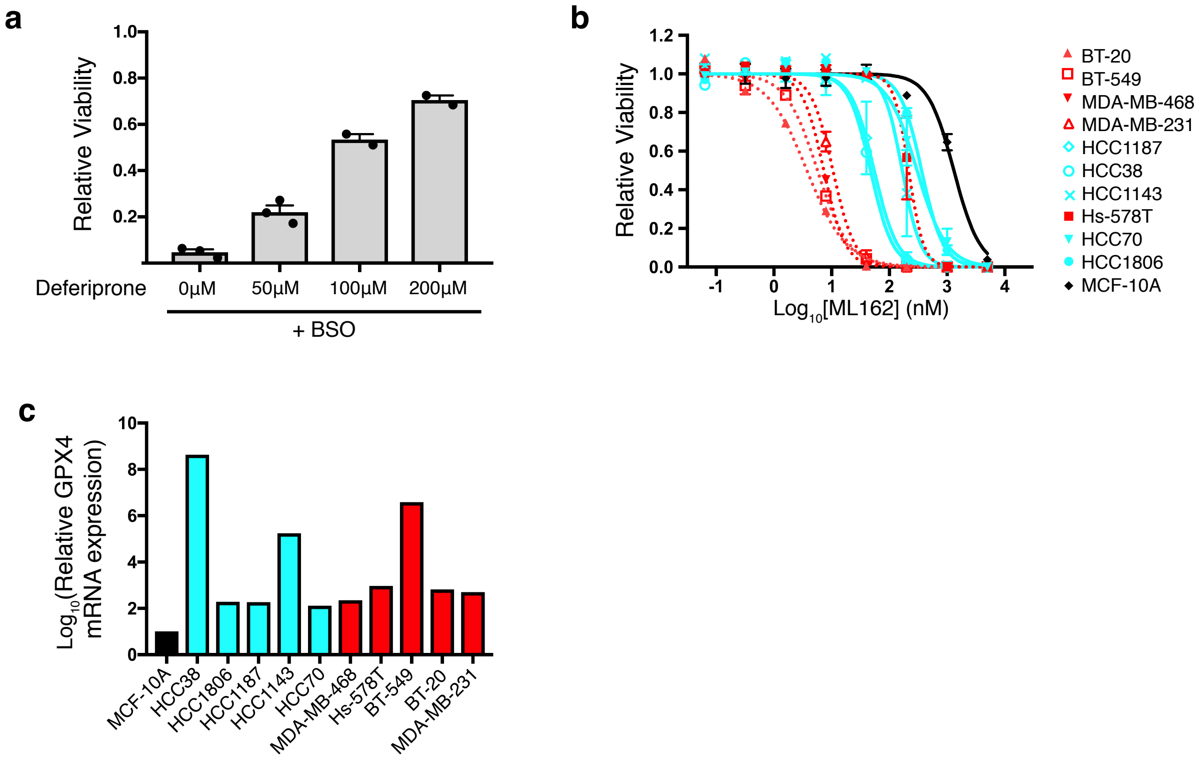

a
Relative Viability
1.0
0.8
0.6
0.4
0.2
Deferiprone
0μM
50μM
100μM
200μM
+ BSO
b
Relative Viability
1.2
1.0
0.8
0.6
0.4
0.2
0.0
-1
0
1
2
3
4
Log10[ML162] (nM)
BT-20
BT-549
MDA-MB-468
MDA-MB-231
HCC1187
HCC38
HCC1143
Hs-578T
HCC70
HCC1806
MCF-10A
c
Log10(Relative GPX4 mRNA expression)
10
8
6
4
2
0
MCF-10A
HCC38
HCC1806
HCC1187
HCC1143
HCC70
MDA-MB-468
Hs-578T
BT-549
BT-20
MDA-MB-231

### Figure S2

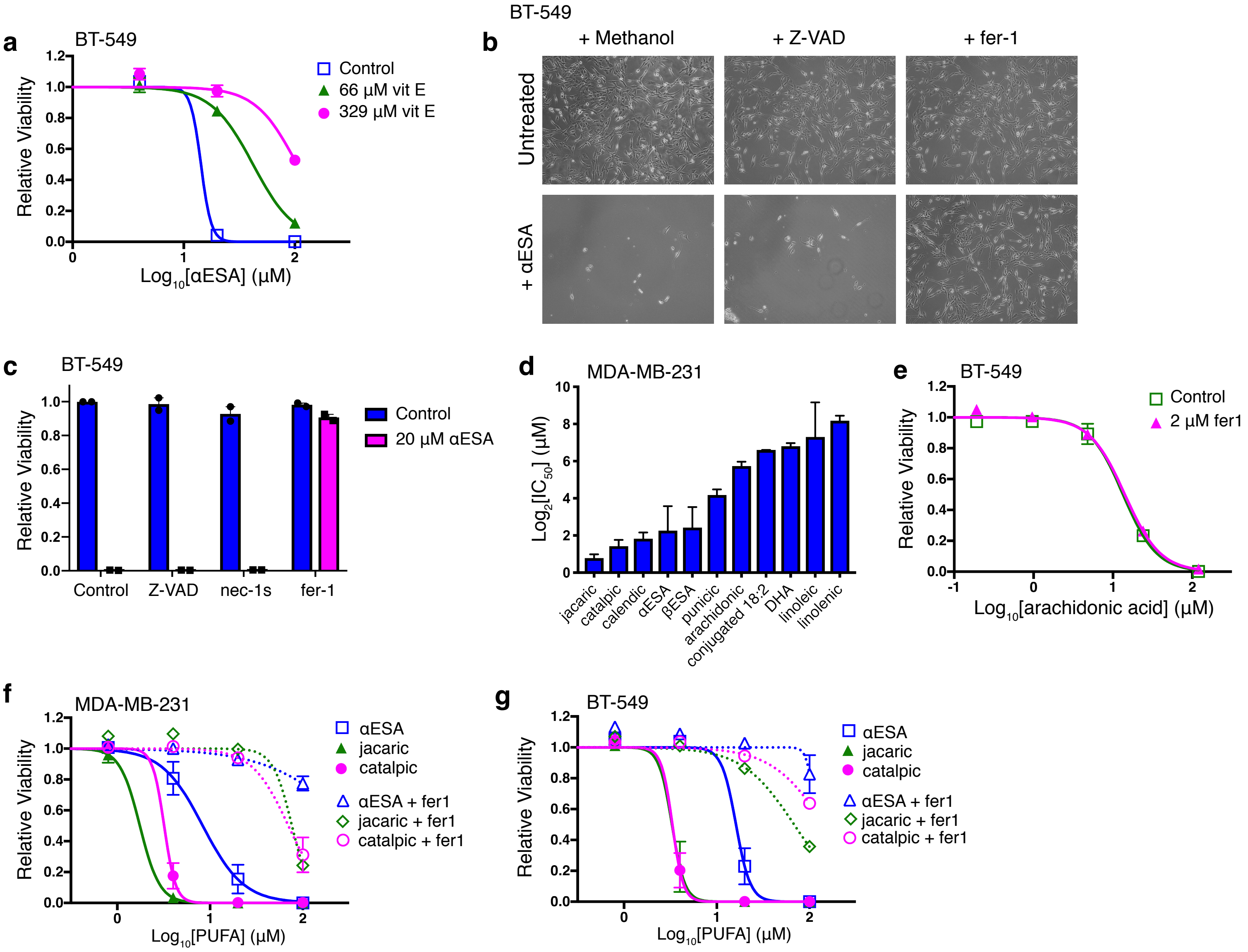
a
BT-549
Relative Viability
Log10[αESA] (μM)
Control
66 μM vit E
329 μM vit E
b
BT-549
+ Methanol
+ Z-VAD
+ fer-1
Untreated
+ αESA
c
BT-549
Relative Viability
Control
Z-VAD
nec-1s
fer-1
Control
20 μM αESA
d
MDA-MB-231
Log2[IC50] (μM)
jacaric
catalpic
calendic
αESA
βESA
punicic
arachidonic
conjugated 18:2
DHA
linoleic
linolenic
e
BT-549
Relative Viability
Log10[arachidonic acid] (μM)
Control
2 μM fer1
f
MDA-MB-231
Relative Viability
Log10[PUFA] (μM)
αESA
jacaric
catalpic
αESA + fer1
jacaric + fer1
catalpic + fer1
g
BT-549
Relative Viability
Log10[PUFA] (μM)
αESA
jacaric
catalpic
αESA + fer1
jacaric + fer1
catalpic + fer1

#### Figure S3

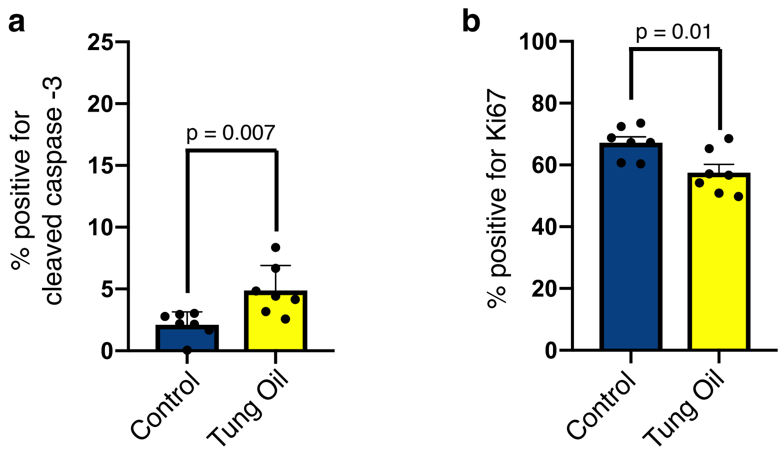

a
% positive for cleaved caspase -3
25
20
15
10
5
0
p = 0.007
Control
Tung Oil
b
% positive for Ki67
100
80
60
40
20
0
p = 0.01
Control
Tung Oil

**Figure S4**

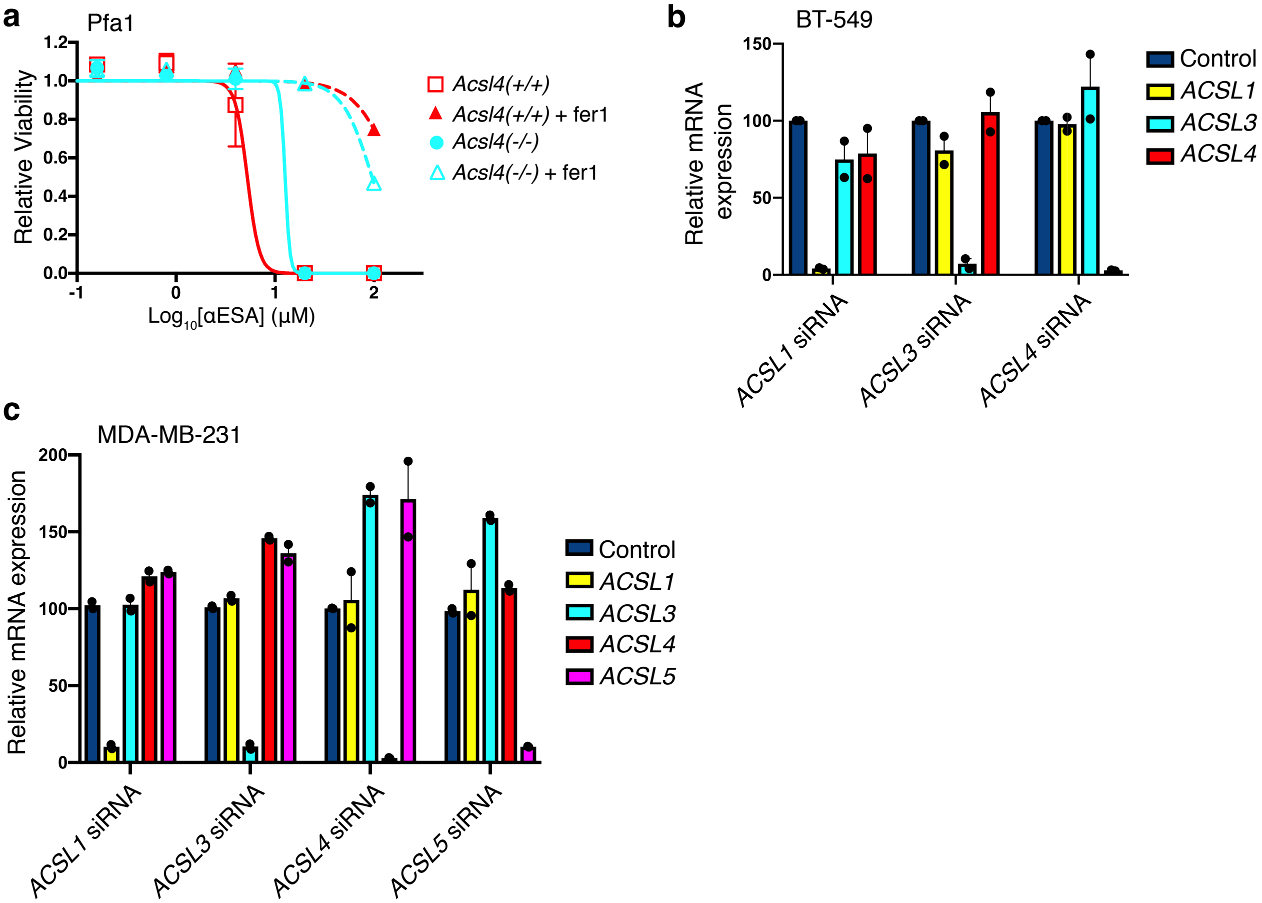

a
Pfa1
Relative Viability
1.2
1.0
0.8
0.6
0.4
0.2
0.0
-1
0
1
2
Log10[αESA] (μM)
Acsl4(+/+)
Acsl4(+/+) + fer1
Acsl4(-/-)
Acsl4(-/-) + fer1
b
BT-549
Relative mRNA expression
150
100
50
0
ACSL1 siRNA
ACSL3 siRNA
ACSL4 siRNA
Control
ACSL1
ACSL3
ACSL4
c
MDA-MB-231
Relative mRNA expression
200
150
100
50
0
ACSL1 siRNA
ACSL3 siRNA
ACSL4 siRNA
ACSL5 siRNA
Control
ACSL1
ACSL3
ACSL4
ACSL5

### Figure S5

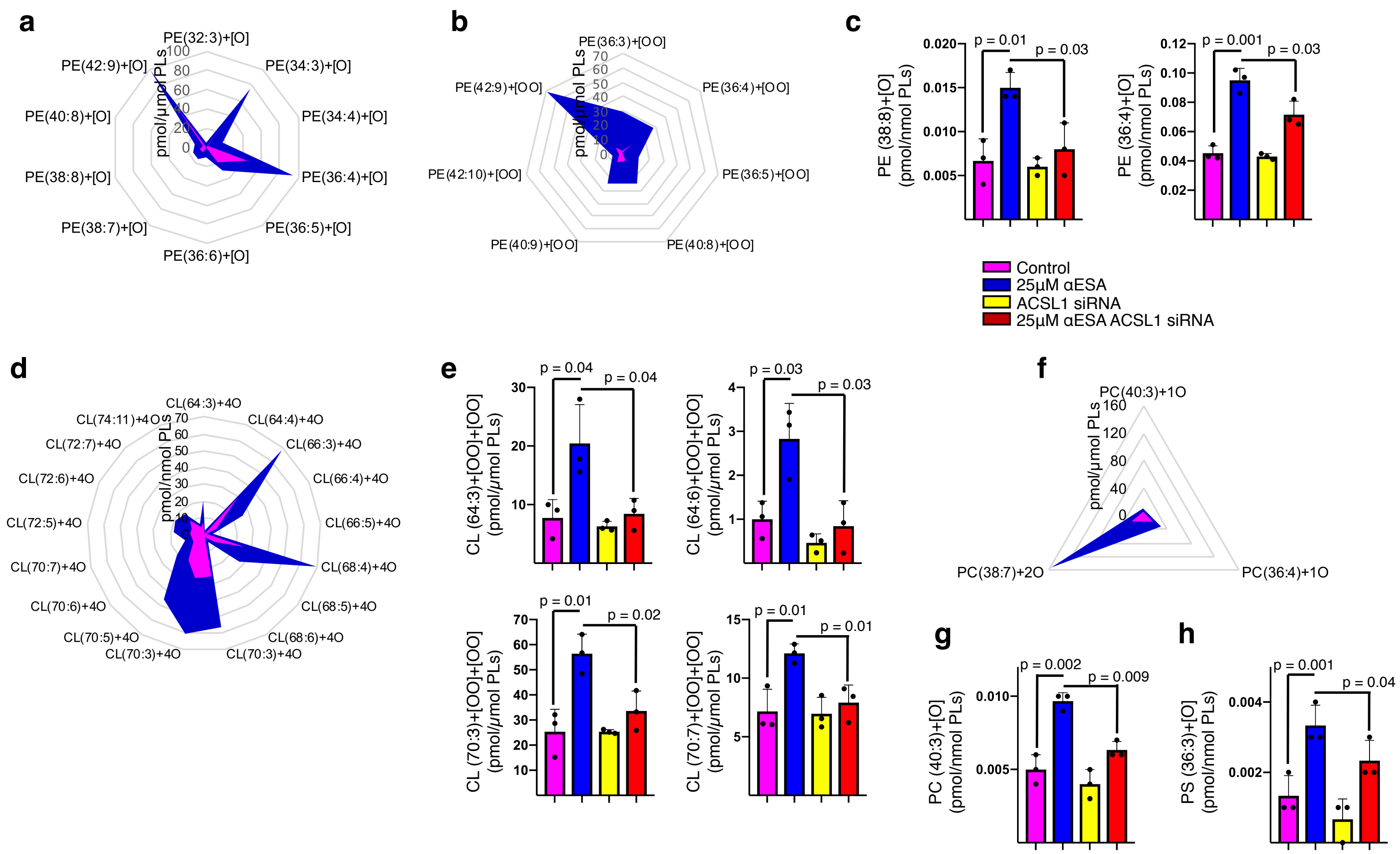
a
PE(32:3)+[O]
PE(34:3)+[O]
PE(34:4)+[O]
PE(36:4)+[O]
PE(36:5)+[O]
PE(36:6)+[O]
PE(38:7)+[O]
PE(38:8)+[O]
PE(40:8)+[O]
PE(42:9)+[O]
pmol/μmol PLs
b
PE(36:3)+[OO]
PE(36:4)+[OO]
PE(36:5)+[OO]
PE(40:8)+[OO]
PE(40:9)+[OO]
PE(42:10)+[OO]
PE(42:9)+[OO]
pmol/μmol PLs
c
PE (38:8)+[O] (pmol/nmol PLs)
p = 0.01
p = 0.03
PE (36:4)+[O] (pmol/nmol PLs)
p = 0.001
p = 0.03
Control
25μM αESA
ACSL1 siRNA
25μM αESA ACSL1 siRNA
d
CL(64:3)+4O
CL(64:4)+4O
CL(66:3)+4O
CL(66:4)+4O
CL(66:5)+4O
CL(68:4)+4O
CL(68:5)+4O
CL(68:6)+4O
CL(70:3)+4O
CL(70:3)+4O
CL(70:5)+4O
CL(70:6)+4O
CL(70:7)+4O
CL(72:5)+4O
CL(72:6)+4O
CL(72:7)+4O
CL(74:11)+4O
pmol/nmol PLs
e
CL (64:3)+[OO]+[OO] (pmol/μmol PLs)
p = 0.04
p = 0.04
CL (64:6)+[OO]+[OO] (pmol/μmol PLs)
p = 0.03
p = 0.03
CL (70:3)+[OO]+[OO] (pmol/μmol PLs)
p = 0.01
p = 0.02
CL (70:7)+[OO]+[OO] (pmol/μmol PLs)
p = 0.01
p = 0.01
f
PC(40:3)+1O
PC(36:4)+1O
PC(38:7)+2O
pmol/μmol PLs
g
PC (40:3)+[O] (pmol/nmol PLs)
p = 0.002
p = 0.009
h
PS (36:3)+[O] (pmol/nmol PLs)
p = 0.001
p = 0.04

### Figure S6

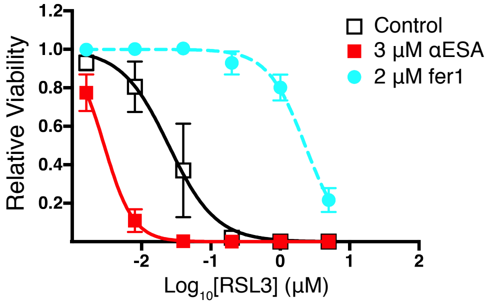
Relative Viability
1.2
1.0
0.8
0.6
0.4
0.2
-2
-1
0
1
2
Log10[RSL3] (μM)
Control
3 μM αESA
2 μM fer1
